# A forest is more than its trees: haplotypes and ancestral recombination graphs

**DOI:** 10.1101/2024.11.30.626138

**Authors:** Halley Fritze, Nathaniel Pope, Jerome Kelleher, Peter Ralph

## Abstract

Foreshadowing haplotype-based methods of the genomics era, it is an old observation that the “junction” between two distinct haplotypes produced by recombination is inherited as a Mendelian marker. In a genealogical context, this recombination-mediated information reflects the persistence of ancestral haplotypes across local genealogical trees in which they do not represent coalescences. We show how these non-coalescing haplotypes (“locally-unary nodes”) may be inserted into ancestral recombination graphs (ARGs), a compact but information-rich data structure describing the genealogical relationships among recombinant sequences. The resulting ARGs are smaller, faster to compute with, and the additional ancestral information that is inserted is nearly always correct where the initial ARG is correct. We provide efficient algorithms to infer locally-unary nodes within existing ARGs, and explore some consequences for ARGs inferred from real data. To do this, we introduce new metrics of agreement and disagreement between ARGs that, unlike previous methods, consider ARGs as describing relationships between haplotypes rather than just a collection of trees.

## 1 Introduction

Ancestral recombination graphs (ARGs) describe how a set of sampled sequences are related to each other at each position of the genome in a recombining species [Brandt et al., 2024, Lewanski et al., 2024, Nielsen et al., 2024, Wong et al., 2024], and there has been significant recent progress on inference through a range of approaches [Rasmussen et al., 2014, Speidel et al., 2019, Kelleher et al., 2019, Zhang et al., 2023, Deng et al., 2024, Gunnarsson et al., 2024]. One way of viewing ARGs is as a sequence of local trees, i.e., the genealogical trees that describe how each portion of the genome was inherited by the focal genomes. This is reflected in methodology of some ARG inference methods and in metrics used to assess inference accuracy, as well as in basic terminology. For instance, the “succinct tree sequence”, introduced by Kelleher et al. [2016], is a common format for describing these inferred ARGs, and is seeing wide use thanks in part to its efficiency and accompanying reliable toolkit, `tskit` [Kelleher et al., 2024, Ralph et al., 2020].

However, an ARG is emphatically not merely a sequence of trees: viewed another way, it describes inheritance relationships between ancestral haplotypes. These two points of view are related because a single haplotype may extend over many local trees; in other words, the internal nodes in the trees are labeled, and many of these labels are shared between adjacent trees [Wong et al., 2024].

Another reason we tend to focus on the trees is that much of our intuition about inference of relationships from genomic data comes from phylogenetics. Indeed, all methods might very roughly be summarized as “more similar sequences are more closely related”. For instance, two sequences that share a derived mutation are probably more closely related over some span of genome surrounding the location where the mutation occurs. It has long been observed that not only mutations but also the “junctions” between distinct haplotypes, if they could be somehow identified, would be inherited as Mendelian markers [Fisher, 1954, Chapman and Thompson, 2003]. In more modern terminology, even in the absence of new mutations, recombination between distinct haplotypes can create a novel haplotype whose relationships and origination time could be inferred.

Haplotype identity has been largely overlooked in the literature on ARG inference – most methods that have been used so far to measure accuracy of inferred ARGs depend only on the sequence of local trees, not on how ancestral haplotypes span across these trees. For instance, Kelleher et al. [2019] and Zhang et al. [2023] compared true and inferred ARGs using average Robinson-Foulds [Robinson and Foulds, 1981] and Kendall-Colijn [Kendall and Colijn, 2016] distances between trees across a regular sequence of genomic positions, using sampled genotypes as labels, while Brandt et al. [2022] compared times to most recent common ancestor between pairs of sampled genomes. Neither is affected by shared haplotype structure – two ARGs could be identical by either measure but imply completely different patterns of haplotype sharing and inheritance. Also, Deng et al. [2021] evaluated agreement of distributions of distances along the genome between tree topology changes, and Zhang et al. [2023] defined a generalization of Robinson-Foulds distance that is the total variation distance between the induced distribution on genotypes; however, neither of these measure the sharing of haplotypes between adjacent trees. An exception is Ignatieva et al. [2024], who compared distributions of haplotype spans in true and inferred ARGs, as well as more sophisticated summaries of edges. The additional information provided by haplotype structure can be important: for instance, haplotypes that extend over many local genealogies “tie together” those genealogies, allowing estimates of times of particular ancestors to be informed by larger portions of the genome on which there are many genealogies.

In this paper, we study various aspects of haplotype identity in ARGs. First, we describe a deterministic algorithm that extends the genomic region spanned by ancestral haplotypes using the principle that intermediate nodes in inheritance paths should remain unchanged when possible. These extended portions of ancestral haplotypes manifest as unary nodes in the local trees. To quantify how accurate the new information is, we define and describe how to compute new measures of (dis-)agreement between ARGs that are motivated by the Robinson-Foulds distance between trees but account for haplotype identity. These measures show that the vast majority of these extended haplotypes are correct if the trees are correct, and that substantial information about haplotypes is contained in these nodes in inferred trees as well.

### 1.1 Motivation and statement of problem

Consider the (small portion of a) hypothetical ARG in Figure 1 A. On the first portion of the genome (lefthand tree), the sample nodes (labeled 0, 1, and 2) coalesce into a small subtree: 1 and 2 find a common ancestor in ancestral node 3, which finds a common ancestor with node 0 in ancestral node 4. On the next portion of the genome (right-hand tree), sample node 2 has a different ancestor. This seems reasonable, and a method that infers trees separately on each portion of the genome could not be expected to produce anything different. However, the example becomes more complicated once we consider what these local genealogies imply about haplotype inheritance. Figure 1 B shows the implied inheritance of haplotypes, with the haplotypes carried by 4 to the left and right of the recombination breakpoint labeled *L* and *R*. Here, sample node 2 has inherited the chunk of haplotype labeled *L* from ancestral node 4 via 3, and the haplotype to the right of this from some other node (and so doesn’t carry haplotype *R*). On the other hand, sample node 1 has inherited *both* haplotypes *L* and *R* from ancestral node 4, but the trees imply that only haplotype *L* is inherited via ancestral node 3. This implies – if taken literally – that there must have been a recombination event at some point between node 1 and node 4 that separated the *L* and *R* haplotypes, and then these two ancestral (and nonoverlapping) haplotypes coalesced together in ancestral node 4. Although this is possible, it seems unlikely – a more parsimonious explanation is depicted in Figure 1 C, in which sample node 1 inherits the entire *LR* haplotype from ancestral node 4 through node 3 (and there is a recombination somewhere between node 3 and node 2). This implies that ancestral node 3 inherits from node 4 on the right-hand tree as well, which is depicted in Figure 1 D – and so node 3 has become unary in this tree. Note that the more parsimonious ARG also includes fewer edges: the three distinct edges 4 → 3, 3 → 1, and 4 → 1 in Figure 1 B have been reduced to the two edges 4 → 3 and 3 → 1 in Figure 1 D.

**Figure 1.**
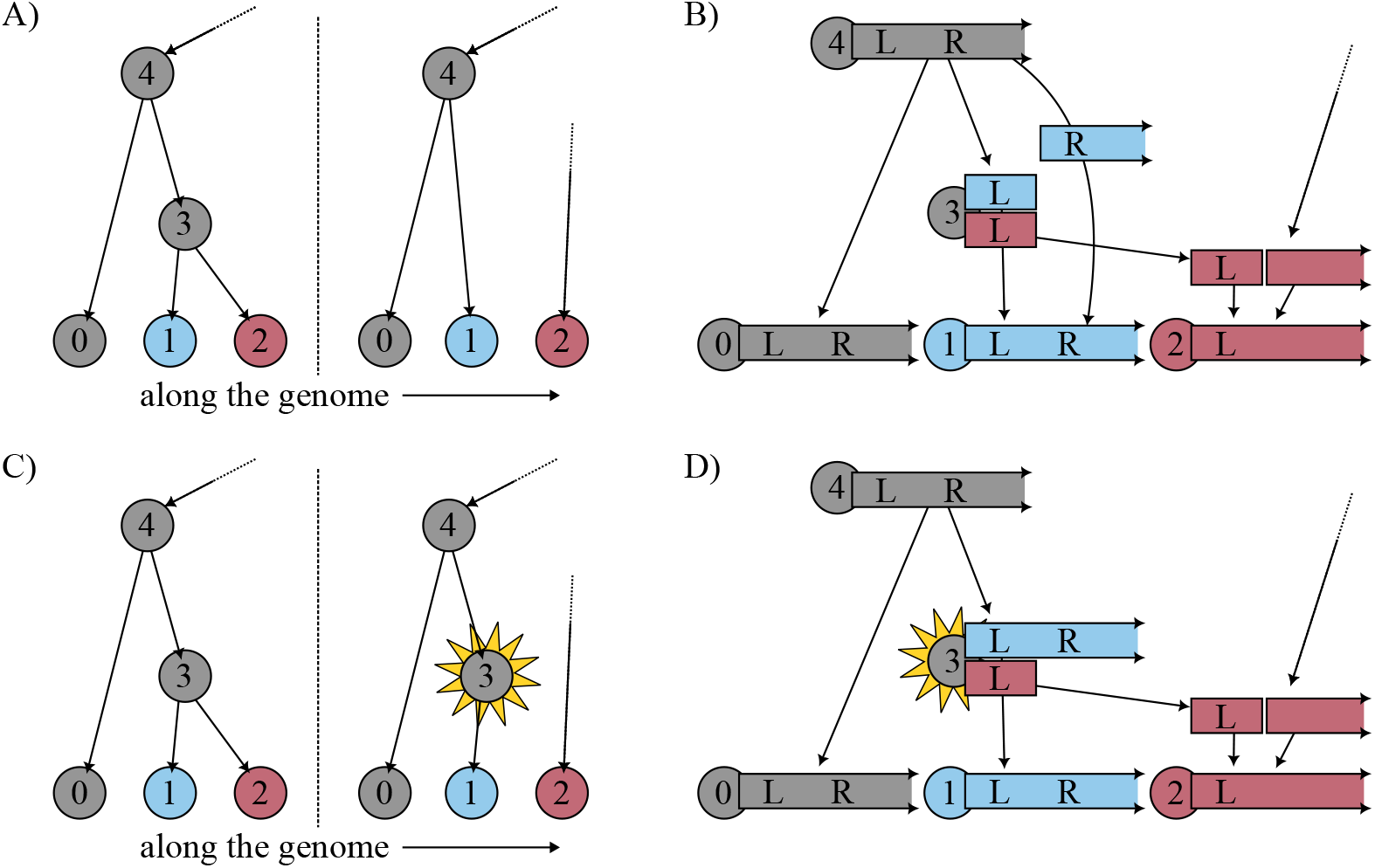
A simple example showing the basic idea (described in more detail in the text): **(A)** a small portion of an ARG without unary nodes; **(B)** the implied inheritance pattern of the two portions of the haplotype carried by ancestral node 4, labeled *L* and *R*; **(C)** local trees with a unary node added, which produces **(D)** a more parsimonious haplotype inheritance pattern (that also includes fewer edges).

So, given the ARG shown in Figure 1 A&B, it should be possible to extend the ancestral haplotype represented by node 3 to obtain the ARG shown in Figure 1 C&D, thus adding additional information to the ARG. This might be surprising, as intuition from phylogenetics suggests we can only infer information about the branching points in the tree, not intermediate (unary) nodes. The goal of this paper is to answer: How can we do this, and how accurate is the resulting inference? See the Discussion for more on what this question assumes, and the connection to parsimony.

## 2 Methods

### 2.1 Notation and terminology

We work with the *succinct tree sequence* representation of ARGs (henceforth, “tree sequence”), to take advantage of the tools available in `tskit` [Kelleher et al., [2024], and our terminology and notation follows Ralph et al. 2020]. For our purposes here, a tree sequence 𝕋 = (*N, E*) contains a set of *nodes N* which represent ancestral segments of genome, and *edges E* which represent relationships between nodes over different regions of the genome. Each node *n* ∈ *N* has a *time t*_*n*_, which is the amount of time in the past that the individual who carried that segment of genome lived. Some nodes are *samples*, meaning that they represent genome sequences available as data. Each edge *e* ∈ *E* describes inheritance between an ancestor and a descendant over a segment of genome [*ℓ*_*e*_, *r*_*e*_). The ancestor and descendant are represented, respectively, by the parent node *p*_*e*_ and child node *c*_*e*_ of the edge and occur at unique times so that 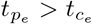. Suppose that the unique elements of the set of left and right edge endpoints are 0 = *a*_0_ *< a*_1_ < … < *a*_*n*_ = *L*, where *L* is the length of the genome. Using this information, one can construct the sequence of trees (*T*_1_, …, *T*_*n*_) that describe how the nodes are related to each other along the genome: each *T*_*k*_ is a tree whose nodes are in *N* and that describes relationships on the half-open interval [*a*_*k*−1_, *a*_*k*_). Nodes represent (portions of) ancestral haplotypes, and so we will use the terms interchangeably. Not all nodes appear in each tree, and we say *n* ∈ *T*_*k*_ for a node *n* if the tree *T*_*k*_ describes at least one parent-child relationship for node *n*.

### 2.2 An algorithm to extend haplotypes

Given a tree sequence, our goal is to identify areas of implied inheritance of haplotypes. Generalizing from Figure 2, we do this by identifying paths of inheritance that are shared across a sequence of local trees but for which some of the intermediate nodes are missing. Concretely, suppose that if in tree *T*_*k*_ there is a chain of inheritance *p* → *u*_1_ → … → *u*_*m*_ → *c* (where *a* → *b* denotes a parent-child relationship) and in tree *T*_*k*+1_ there is a chain of inheritance *p* → *v*_1_ → … → *v*_*n*_ → *c*, where 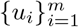 and 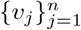 are disjoint. This situation implies that *c* inherited from *p* over the entire interval [*a*_*k*−1_, *a*_*k*+1_), so it seems reasonable to assume that *c* has inherited from *p along the same path* for that entire interval. In other words, the intermediate nodes {*u*_*i*_} should also lie on the path from *c* to *p* in tree *T*_*k*+1_, and conversely the nodes {*v*_*j*_} should lie on that path in tree *T*_*k*_. For instance, in Figure 1 take *p* = 4 and *c* = 1, so that *u*_1_ = 3 (and *m* = 1) and there are no *v* (so *n* = 0); and so as shown in Figure 1 B&D, we might extend node 3 over this entire segment. Of course, this does not always make sense – for instance, if *u*_*i*_ is already represented somewhere else in *T*_*k*+1_, or if 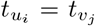, for some *i* and *j*. So, we restrict our attention to pairs of such paths in adjacent trees for which *u*_*i*_ ∉ *T*_*k*+1_ for all 1 ≤ *i* ≤ *m, v*_*j*_ ∉ *T*_*k*_ for all 1 ≤ *j* ≤ *n*, and the times of the nodes {*u*_*i*_} and {*v*_*j*_} are unique, meaning no node in *u*_*i*_ has the same time as a node in *v*_*j*_. Call a pair of such paths *mergeable*. So, the goal of our algorithm is to iterate over trees, identify mergeable pairs of paths, and then extend the nodes {*u*_*i*_} to *T*_*k*+1_. (We also extend {*v*_*j*_} to *T*_*k*_, but on a backwards pass.)

**Figure 2.**
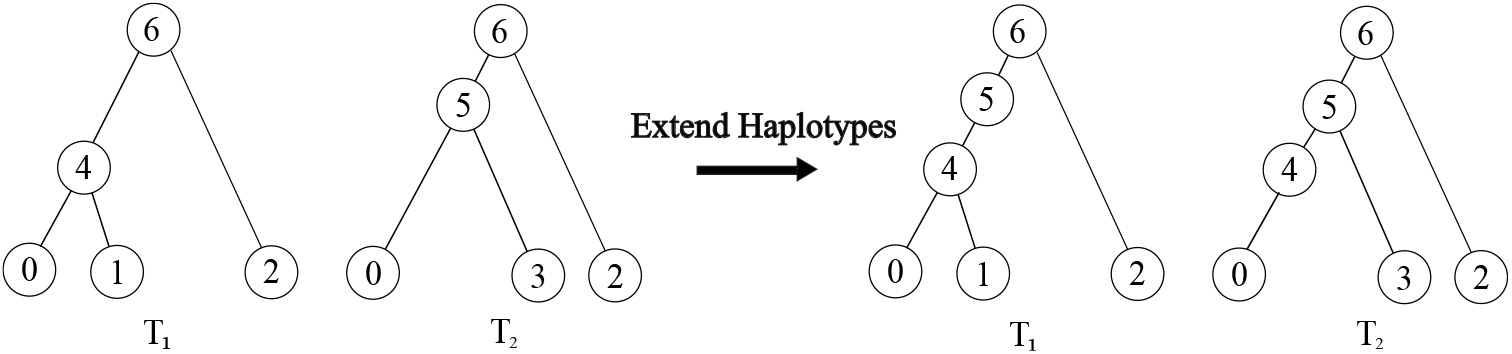
A visualization of the *extend haplotypes* method. In both trees *T*_1_ and *T*_2_, node 0 inherits from node 6, the root: *T*_1_ contains the path 6 → 4 → 0 while *T*_2_ has path 6 → 5 → 0. The intermediate nodes 4 and 5 do not appear in *T*_2_ and *T*_1_ respectively, and so the paths are *mergeable*. The “extend haplotypes” method joins these two paths, inserting the merged path 6 → 5 → 4 → 0 into both *T*_1_ and *T*_2_.

An efficient algorithm to do this is described in Algorithm 1. The algorithm considers each tree transition from *T*_*k*_ to *T*_*k*+1_ in turn, updating its internal state (which includes possibly modifying *T*_*k*+1_) as it goes. Suppose we are at the transition from tree *T*_*k*_ to tree *T*_*k*+1_, which is done by first removing a set of edges *O* and then adding another set of edges *I. O* defines a sub-forest *F*_*O*_ of *T*_*k*_, and *I* defines a sub-forest *F*_*I*_ of *T*_*k*+1_. In Figure 1 (in the portion of the tree shown), the removed edges that make up *F*_*O*_ are 4 → 3, 3 → 1, and 3 → 2, and the added edges that make up *F*_*I*_ are 4 → 1 and the new edge leading to 2. The key step in the algorithm is to determine whether the pair of paths that terminate in a given node in two adjacent trees are mergeable. The algorithm we use to do this is given as Algorithm 2, and works as follows. If a pair of paths is mergeable, then the edges of the two paths must lie in *O* and *I*, respectively. Suppose an edge in *I* has child *c*. To see if *c* is the base of a pair of mergeable paths, the algorithm traverses up from *c* in both *F*_*O*_ and *F*_*I*_ ; the variables *y*_*i*_ and *y*_*o*_ indicate whether the next node upwards in the traversals (i.e., the next older node, as recall *t*_*n*_ is amount of time in the past that *n* lived) is in *F*_*I*_ or *F*_*O*_. The traversals terminate if a node in the other tree is found (i.e., if the node traversed in *F*_*O*_ is in *T*_*k*+1_ or if the node traversed in *F*_*I*_ is in *T*_*k*_) or if a pair of traversed nodes have the same time. If these two traversals end in the same node *p*, the paths are mergeable. Iterating over all edges in *O* will thus find all mergeable pairs of paths. There is often more than one pair of mergeable paths in a tree transition; so, the algorithm merges pairs of mergeable paths, starting with pairs that add the smallest number of new edges, until no more are found.

Note that the algorithm could be applied to an undated ARG, with some adjustments. The algorithm only uses node time for two reasons: convenience, when iterating jointly up the two paths; and to avoid illegal conditions: if we tried to merge two paths on which some *u*_*i*_ had the same time as some *v*_*j*_, then we would have a parent and child with the same time, which is not allowed.

Algorithm 1 simplifies the full algorithm implemented in software in several ways for the sake of clarity – for instance, the bookkeeping required to keep track of *T*_*k*_ and *T*_*k*+1_ is omitted. Furthermore, as described the algorithm does one left-to-right pass over the tree sequence; in practice we do repeated passes in both directions until no changes can be made. We require repeated passes because the additional structure added by a given path of the algorithm may introduce more mergeable paths. Most of these cases seem to occur for paths with large time differences between the parent and child nodes, or when many nodes in a path are coalescent. The number of required passes is in practice small: see empirical results below.

The main step that is omitted is a description of the `Merge` operation, which performs the actual extending of haplotypes. This algorithm is essentially the same as `MergeNum` in Algorithm 2, except with additional bookkeeping. Roughly speaking, the algorithm traverses up from the shared base node *c*, doing the appropriate operations to insert the nodes along the path found in *T*_*k*_ into the path in *T*_*k*+1_. To do this, some edges that end at *a*_*k*_ will be extended to end at *a*_*k*+1_; some edges that begin at *a*_*k*_ will be postponed to begin at *a*_*k*+1_, and some entirely new edges may be added, as in Figure 2. Furthermore, the forests *F*_*O*_ and *F*_*I*_ (and hence *T*_*k*+1_) need to be updated.

#### Algorithm 1

Extend haplotypes. Given an ARG 𝕋 with *N* trees *T*_1_, …, *T*_*N*_ for which edges *I*_*k*_ are added to transition from *T*_*k*_ to *T*_*k*+1_, identify and merge all mergeable paths (see text). Each child node *c*_*e*_ of each removed edge *e* is checked to see if it is at the base of two mergeable paths; paths that add fewer new edges are merged first because `MergeNum` returns the number of new edges required to merge the two paths, and *M* is always less than or equal to the minimum number of new edges across all nodes in the current tree transition.

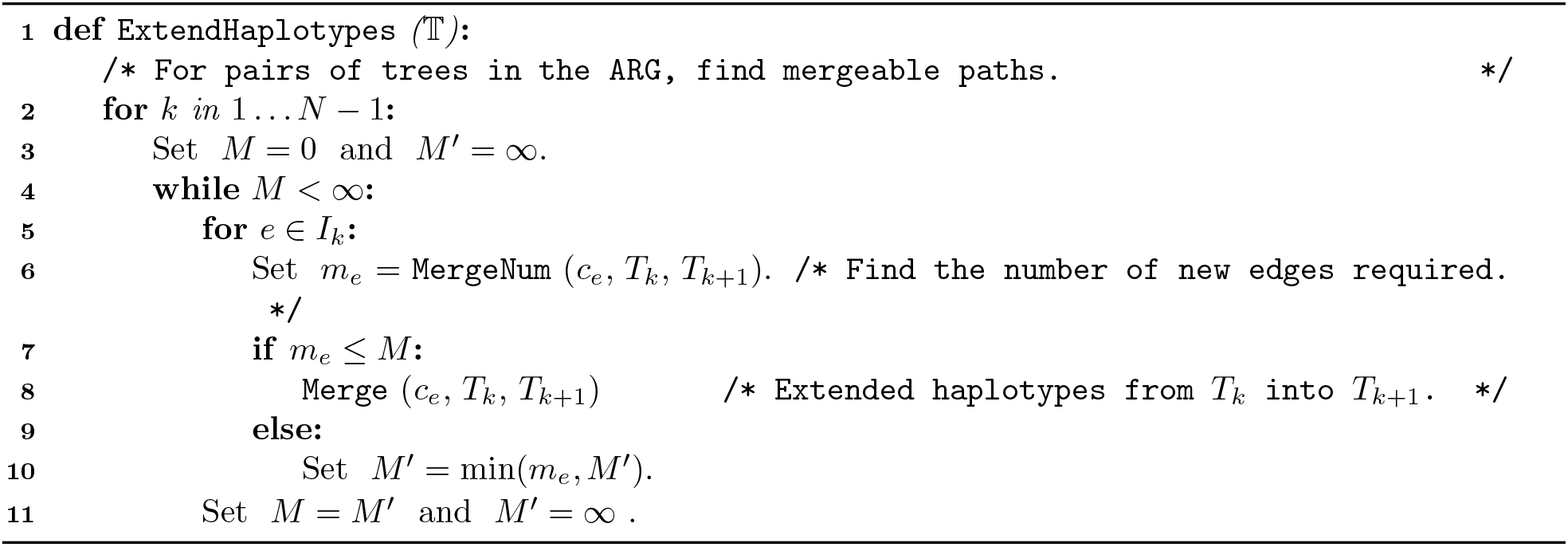

Because the algorithm needs to take multiple passes over the tree sequence in each direction, an important practical question for this algorithm is: how many passes do we need to do? The algorithm is monotone (spans of ancestral nodes only increase), so it is guaranteed to terminate in a finite number of passes, but it is also not hard to construct pathological cases that require an arbitrary number of passes. However, experimentation suggests that in practice at most five iterations are needed before the algorithm terminates. Indeed, for even large sequences Table S1 shows that 99% of all changes to an ARG occur after the first iteration, with the algorithm always completing after four iterations.

#### Algorithm 2

Given a node *c*, trees *T*_*O*_ and *T*_*I*_, and sub-forests *F*_*O*_ and *F*_*I*_ such that removing *F*_*O*_ and adding *F*_*I*_ turns *T*_*O*_ into *T*_*I*_, check to see if the paths upwards from *c* in *T*_*O*_ and *T*_*I*_ are mergeable. If the paths are mergeable then this returns the number of new edges that would be added by extending the path from *T*_*O*_ to *T*_*I*_ ; otherwise, this returns *∞*. Let *P*_*O*_[*n*] and *P*_*I*_ [*n*] be the parents of node *n* in the set of edges to be removed and added, respectively (i.e., in *F*_*O*_ and *F*_*I*_); these are NULL if *n* has no parent, and we take *t*[NULL] = *∞*. The variable *m*_*e*_ will record the number of new edges to be added, and *m* will record the number of extended haplotypes. Note that if no haplotypes can be extended (*m* = 0), we return *m*_*e*_ as *∞* so that the path is not merged in Algorithm 1.

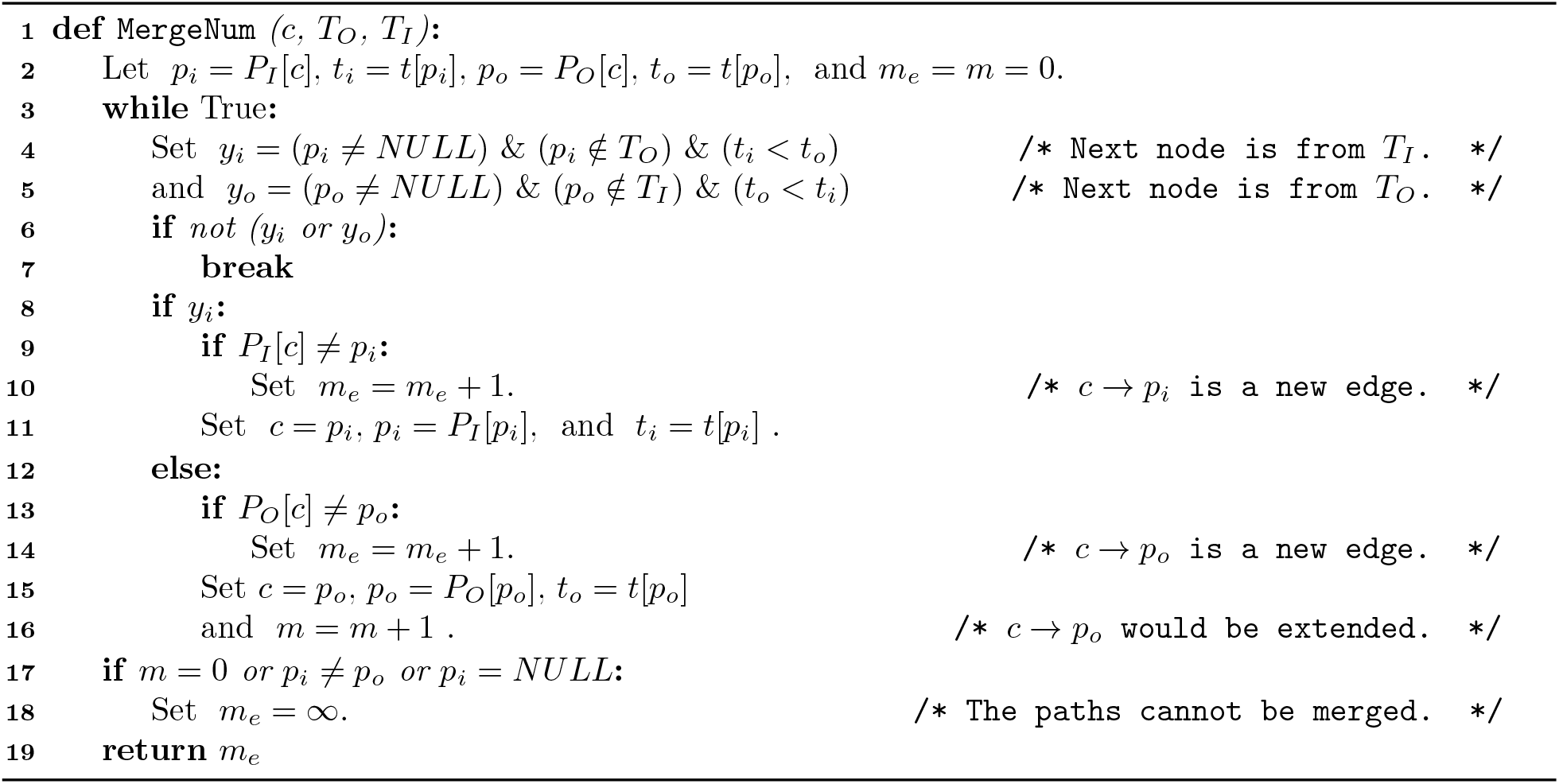

### 2.3 Dissimilarity between ARGs

If we begin with a tree sequence containing unary nodes, it is straightforward to remove the portions of each node’s span on which it is unary, extend haplotypes (Algorithm 1), and quantify how much node span was correctly or incorrectly added. However, we are also interested in whether extending haplotypes improves *inferred* ARGs. Since we are not aware of any current methods for measuring (dis)agreement between ARGs that take into account haplotype identity, we define a measure of *matched span* to quantify this. The method (including Equation 1 - Equation 5) is implemented in the `tscompare` package.

It is helpful to first describe what we compute in the simple case. We will first simulate tree sequences where nodes that are unary in local trees between coalescent haplotypes are retained. Then, each node is present in both tree sequences, and we can quantify, for each node, how much of their span is correct or incorrect by comparing to the original, true tree sequence.

Now suppose that instead of comparing two tree sequences with the same set of nodes, we wish to compare two tree sequences for which we know the sample nodes are the same but are otherwise unclear as to the equivalency of nodes across sequences. (For instance, with a simulated tree sequence and one inferred from its genotypes; nodes in the former represent actual ancestral haplotypes, and in the latter represent hypothetical ancestors which may or may not resemble the truth.) Call the two tree sequences 𝕋_1_ and 𝕋_2_, which should have the same genome length and the same set of sample nodes; in what follows we think of 𝕋_2_ as the true ARG and 𝕋_1_ as an inferred ARG. We would like to measure (a) how much of 𝕋_1_ is found in 𝕋_2_; (b) how much of 𝕋_2_ is found in 𝕋_1_; and (c) how much of 𝕋_1_ is *not* found in 𝕋_2_. (Think of these three quantities as the sizes of two relative intersections and difference between the tree sequences, thought of vaguely as sets; so note in particular that switching the roles of 𝕋_1_ and 𝕋_2_ gets an different set of quantities.) Roughly speaking, we first identify matching nodes as those whose sets of descendant samples agree for the largest span along the genome, and then compute for how much of their spans do their descendant samples agree (or not). An example of our method is illustrated in Figure 3.

**Figure 3.**
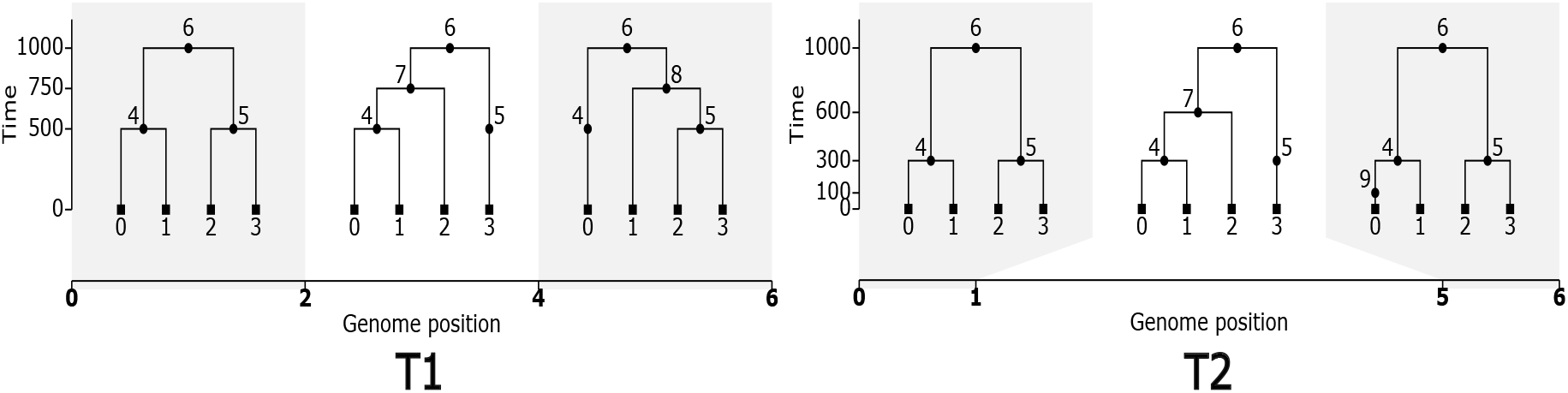
For two tree sequences *T* 1 and *T* 2 the *matched span*, 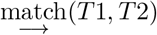, matches nodes in *T* 1 to nodes in *T* 2 based on identical sample sets. We first can compute the total span of nodes in *T* 1 and *T* 2 as ∥*T* 1∥ = 46 and ∥*T* 2 ∥= 47. First, we always require that sample nodes match to their identical counterpart. Then we match intermediate nodes by comparing their descendent samples across pairs of local trees over the genome. Here we have distinct local tree pairings on the genome segments: [0, 1), [1, 2), [2, 4), [4, 5), and [5, 6). Node 4 has no match on [4, 5), matches with node 9 on [5, 6), and matches with node 4 in *T* 2 everywhere else. Thus the maximal mapping for 4 should be to itself since *m*(4, 4) *> m*(4, 9). Nodes 5, 6, and 7 all match with their identical counterpart. Lastly, node 8 has no match in *T* 2 as there are no nodes in *T* 2 with sample set {1, 2, 3}. This makes the matched span 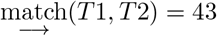 and 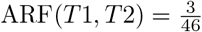. Given the above matching, the inverse matching will match nodes 0 through 7, and node 9 has no match since its only possible match (4 ∈ *T* 1) was not the best match from *T* 1 → *T* 2. This means the inverse matched span 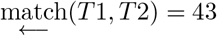 and 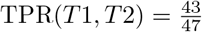.

The method works as follows. To simplify notation suppose that the two tree sequences have the same set of breakpoints between trees, so that 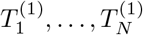 are the trees in 𝕋_1_ and 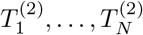 are the trees in 𝕋_2_. For a node *n* and tree *T* let *S*(*T, n*) denote the set of samples that inherit from *n* in *T*, and for a pair of nodes *n*_1_ and *n*_2_ with *n*_1_ in 𝕋_1_ and *n*_2_ in 𝕋_2_, define

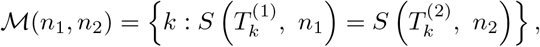

to be the indices of all trees where *n*_1_ and *n*_2_ are ancestral to the same sample set in both ARGs, and

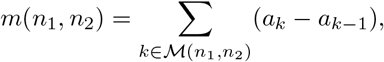

which is the total span over which the samples below *n*_1_ in 𝕋_1_ matches the samples below *n*_2_ in 𝕋_2_. The *matched span* of 𝕋_1_ in 𝕋_2_ is then defined to be

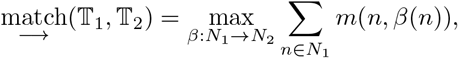

where the maximum is over all mappings *β* of nodes in 𝕋_1_ to nodes in 𝕋_2_, and we require that samples in 𝕋_1_ are mapped to samples in 𝕋_2_. (Note that multiple nodes in 𝕋_1_ may be mapped to the same node in 𝕋_2_, and that some nodes in 𝕋_2_ may not be mapped to by any nodes in 𝕋_1_.) Since the maximum is independent over nodes, we may define for each node *n*_1_ ∈ 𝕋_1_ its *best matching node* in 𝕋_2_ as

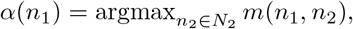

so that

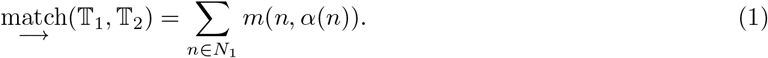

If the best-matching node is not unique, we define *α*(*n*_1_) to be the node in *T*_2_ out of those maximizing *m*(*n, n*) that minimizes 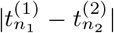 (and if *this* is not unique, we pick an arbitrary one) – however, this potential ambiguity does not affect the definition of 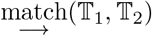. Let *s*(𝕋, *n*) denote the total span that node *n* is present in the local trees,

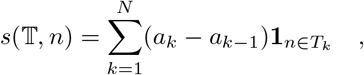

where 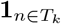 is an indicator (i.e., it is 1 if *n* ∈ *T*_*k*_ and 0 otherwise), and let 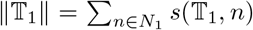 be the total span of all nodes in 𝕋_1_. We then define the *non-matched span* of 𝕋_1_ in 𝕋_2_ by

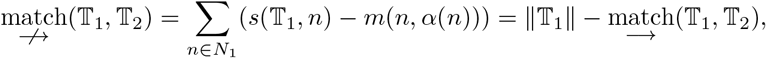

which is the total span for all nodes in 𝕋_1_ over which their descendant samples do *not* match those of their best match in 𝕋_2_. Contrarily, given a matching *α* : 𝕋_1_ → 𝕋_2_, we want to quantify how much of 𝕋_2_ is represented in 𝕋_1_. To do this, we define the *inverse matched span* of 𝕋_1_ in 𝕋_2_ as

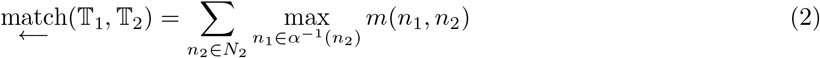

where *α*^−1^(*n*_2_) is the set of all nodes *n*_1_ ∈ 𝕋_1_ whose best match is *n*_2_. This differs from the matched span of 𝕋_2_ in 𝕋_1_ because there may be more than one node in 𝕋_1_ that is mapped to the same node in 𝕋_2_ – so, if nodes *n*_1_ and 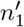 are both mapped by *α* to the same node *n*_2_, then both count towards 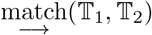, but only the better match counts towards 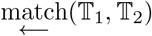.

A common measure of disagreement between ARGs, first proposed by Kuhner and Yamato [2015], is to use a weighted average Robinson-Foulds (RF) distance. This could be computed in a very similar way: instead of *m*(*n, α*(*n*)) define

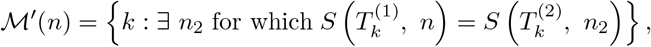

the indices of all trees on which there is *some* node in 𝕋_2_ whose set of descendant samples matches those of *n*, and

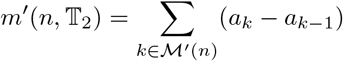

the total span over which *n* finds a match. Then the average RF distance (averaged over locations in the genome) is

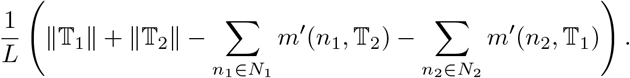

(This holds because the average number of nodes per local tree in 𝕋_1_ is ∥𝕋_1_∥ */L*, so the difference gives the average number of non-matching nodes.) In other words, we require a node in 𝕋_1_ to match the *same* node in 𝕋_2_ across all trees, but average RF distance allows a different node to match on each tree. The other differences are that *average* RF distance normalizes by sequence length, and is symmetrized. The RF distance between two trees was defined by Robinson and Foulds [1981] to be the minimum number of branch contraction/expansion operations needed to move from one tree to the other (which they then show is equal to the number of branches that induce different splits on the labels). A similar metric on ARGs could be defined using the subgraph-prune-and-regraft moves used by Deng et al. [2024].

The matched span, 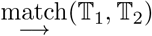, measures agreement between *topologies*, but not times. If the ARG is dated [e.g., as in Wohns et al., 2022, Deng et al., 2024], we can naturally use the “best match” *α* to also compare times. Empirically, dating error seems to be more or less homoskedastic on a log scale, so we recommend using the weighted root-mean-squared error of log(times), computed as

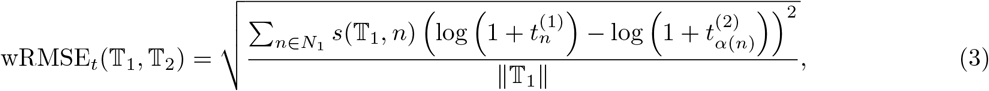

where the transformation is *t* ⟼ log(1 + *t*) to avoid log(0). The mean is computed weighting by node span, so that a dating error is more impactful for a node with a longer span.

The implementation of this method in `tscompare` additionally produces relative values: the ARG RF value (non-matched span relative to 𝕋_1_) and true proportion represented (inverse matched span relative to 𝕋_2_). The ARG RF (which we call “ARF”) is defined to be the non-matched span proportional to the total span of nodes in 𝕋_1_,

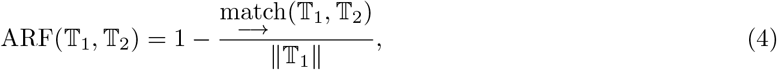

and so if 𝕋_2_ represents the truth, is analogous to a false positive rate. The *true proportion represented* (TPR) is the inverse matched span between two trees relative to the total span of nodes in 𝕋_2_,

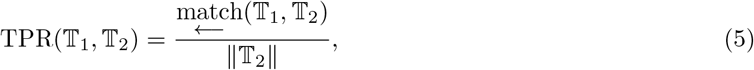

and is analogous to statistical power. The outputs framed as proportions relative to one of the given tree sequences is more easily understood for comparing pairs of tree sequences than the original matched span and inverse matched span, whose units are length of spans. We give an example computing the ARF and TPR between two small tree sequences in Figure 3.

Additionally, let’s compute ARF(*T* 2, *T* 1) and TPR(*T* 2, *T* 1). Again, the sample nodes, root 6, and nodes 4, 5, and 7 will match to their identical counterpart and have matched nodes spans 6 (for each sample), 6, 4, 4, and 2, respectively. Node 9 matches to node 4 with a matched node span of 1. Thus,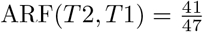. The inverse matched span will match nodes 5, 6, 7 and sample nodes to their identical counterparts. Node 8 in *T* 1 had no match from *T* 2, so it has no inverse match, while node 4 had two best matches from *T* 2: nodes 4 and 9. Since 4 in *T* 1 shares more span with the same-numbered node in *T* 2 than with 9, the inverse match for 4 in *T* 1 is node 4 in *T* 2. Therefore, 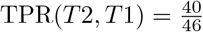.

#### Metrics on ARGs

Neither the matched span or non-matched span of 𝕋_1_ in 𝕋_2_ are metrics in the mathematical sense (i.e., symmetric, nonzero distance between distinct points, and satisfying the triangle inequality). This is by design: in practice it is not possible to infer all aspects of the true ancestry of a set of samples (i.e., all their genetic ancestors who ever lived), and so we wanted to quantify “How much of the true relationships does this ARG represent?” Indeed, even a symmetrized version of non-matched span is not a metric: for instance, if 𝕋_2_ is produced from 𝕋_1_ by adding a new, entirely unary node, then ARF(𝕋_1_, 𝕋_2_) = ARF(𝕋_2_, 𝕋_1_) = 0. This is again by design: in general, edges in ARGs represent transmission of genetic material through many ancestors, and so addition of entirely unary ancestors on an edge is (arguably) not wrong. (Note that our metrics do distinguish these two ARGs: in this example TPR(𝕋_1_, 𝕋_2_) < 1.)

A related (but not identical) way to construct a metric on (undated) ARGs modifies the definitions above so that *α* is a bijection. To do this, write *N*_1_ and *N*_2_ for the nodes of 𝕋_1_ and 𝕋_2_ respectively, and suppose that *φ* is a bijection from *N*_1_ to *N*_2_. Since we want to allow nodes to remain unpaired, for notational convenience suppose that both *N*_1_ and *N*_2_ have appended an arbitrary number of “unrelated” nodes (since these are not ancestral to any samples they do not affect the metrics). Then, define *d*_*φ*_(𝕋_1_, 𝕋_2_) to be the total span of nodes *n*_1_ ∈ *N*_1_ on which *n*_1_ and *φ*(*n*_1_) do not match (i.e., are not ancestral to the same set of samples), plus the total span for nodes *n*_2_ ∈ *N*_2_ on which *n*_2_ and *φ*^−1^(*n*_2_) do not match. The quantity min_*φ*_ *d*_*φ*_(𝕋_1_, 𝕋_2_), where the minimum is over bijections that preserve the partial ordering induced by the ARG, defines a metric on undated ARGs. (Actually, as written it defines a metric on the equivalence classes of undated ARGs that are obtained by removing any structure not ancestral to any samples.) To see this, note that relative ordering and descendant sample sets uniquely determine the (equivalence class of an) ARG, and that the triangle inequality is satisfied because 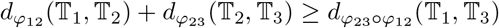.

This metric, min_*φ*_ *d*_*φ*_(𝕋_1_, 𝕋_2_), differs from the definitions above because we require a strict one-to-one match between nodes in 𝕋_1_ and 𝕋_2_ (i.e., *φ* is bijective, while there may be more than one node in 𝕋_1_ that *α* matches to the same node in 𝕋_2_). We did not require this property when defining non-matched span above for two reasons: first, the arguable non-wrongness of additional unary nodes, and second, requiring a bijection makes computation much more difficult.

The RF distance [Robinson and Foulds, 1981] essentially counts the number of differing branches between two trees; the averaged RF distance [Kuhner and Yamato, 2015] averages this distance across local trees, weighted by span along the genome. The method we present here for measuring dissimilarity between topologies of ARGs is a straightforward generalization that takes into account span along the genome of inferred ancestral haplotypes (and separates the metric into two pieces). However, the RF metric has many undesirable properties – for instance, moving a single tip can result in a tree with maximum distance to the original – and there is a substantial literature giving more robust generalizations [reviewed by Llabrés et al., 2021]. Many of these generalizations [e.g., Böcker et al., 2013] relax the requirement that the match between subtended sample sets be exact, and weight matches in some way by the size of the dissimilarity. We considered such definitions as well, but kept to the simple case for computational tractability – the generalization of Böcker et al. [2013] is NP-hard to compute, even for a single tree. In the ARG literature, Zhang et al. [2023] defines a metric (called “ARG total variation distance”) that includes branch lengths, in a way similar to Robinson and Foulds [1979] and Kuhner and Felsenstein [1994]; however, it is still applied to ARGs as an average over local trees, without enforcement of identity across haplotypes; it would be useful to extend our dissimilarity to include branch lengths.

### 2.4 Simulations

Our method for extending haplotypes is applicable to any ARG. However its accuracy depends on the overall structure of the ARG it is applied to. Thus to understand how well our methods can infer ancestral haplotypes we work with ARGs simulated across a range of parameter values. To do this, we simulate ARGs containing full haplotypes using Hudson’s algorithm as implemented in `msprime` [Kelleher et al., 2016, Baumdicker et al., 2021], with the `coalescing_segments only` option set to False. Although `msprime` simulates many events that do not create a coalescence in some local tree, by default it only outputs information for nodes which contain a coalescence (i.e., are the MRCA of some pair of samples at some point on the genome). Furthermore, by default it only outputs those segments of the genome on which there is a coalescence. Said another way, by default all ancestral nodes in an ARG output by `msprime` are the MRCA of some pair of samples at every point in the genome on which they are represented. However, here we are interested in those segments of genome on which the nodes are *not* coalescent; i.e., where they are unary in the local trees. Setting `coalescing_segments only` to False includes just this information: any ancestral segments for which these coalescent nodes are ancestral to any samples – so, the unary portions of their spans as well. However, this includes more information than we want: we hope to recover those portions of ancestral haplotypes on which the nodes are unary, but adjacent to a region of the genome where the node is not unary. For instance, if a lineage carrying an ancestral segment of genome that spans [*a, c*) coalesces with another spanning [*b, d*), with *a* < *b* < *c* < *d*, then the resulting node is only coalescent on [*b, c*) but we hope using this algorithm to extend the node’s span to [*a, b*) and [*c, d*) (on which the node is unary). However, following this example, the first lineage might also carry a segment [*x, y*) that is disjoint from the segment [*a, d*). We call these segments “isolated non-coalescent segments”; they have also been called “trapped unary spans” [by Wong et al., 2024]. Such isolated segments will not be recovered by our algorithm, and would likely be unrecoverable by any other method. So, after simulation, we first remove these isolated, non-coalescent segments. To give an idea of what proportion of the full spans of ancestral nodes these isolated non-coalescent segments represent, a simulation of 1000 samples with genome length 5 *×* 10^7^, recombination rate 10^−8^, and population size 10^4^ has about half the total span of all nodes in isolated, non-coalescent segments. For more discussion of these segments, see Baumdicker et al. [2021].

We used simulations of several scenarios. To include the effects of heterogeneous recombination rate, in some we used stdpopsim [Adrion et al., 2020] to simulate chromosome 1 of *Canis familiaris* using the CanFam3 genetic map from Campbell et al. [2016]. *“Constant dog”* simulations simulated this chromosome in a population of (constant) size 10^4^. *“Expanding dog”* simulations were similar, but used a discrete-time Wright–Fisher model to simulate a small population of 100 that expanded to 1,000 individuals ten generations ago, which then doubled every generation to reach 512,000 individuals. All jobs for which runtime was recorded were executed on an Intel Xeon Gold 6148 processor. Additionally, using our matched span methods, we compare accuracy between sequences modified from a “true” ARG, (see Figure 6). These ARGs were simulated with an effective population size of 10,000 and recombination rate of 10^−8^ with between 10 and 1,000 samples and a genome length between 10^6^ to 5 *×* 10^7^.

## 3 Results

We next present the results of various experiments with inferred ARGs. First, we quantify the reduction in number of edges and resulting computational speedups. Next, to validate the principle of the algorithm, we describe accuracy of inferred additional edge spans when applied to true ARGs with unary spans removed. Finally, we explore implications for accuracy with inferred ARGs.

### 3.1 Tree sequence compression and computation

In the simple example in Figure 1, extending haplotypes replaces three edges (0 → 3 and 3 → 4 on the left tree, and 0 → 4 on the right tree) by two edges (0 → 3 and 3 → 4 on both trees). If all edge endpoints were unique, then we’d expect *every* edge to be extendable on one of its ends (except those pendant to the root and some of those adjacent to chromosome ends), leading to a reduction in number of edges by almost exactly one third. Experiments with an earlier version of the algorithm showed that if we only extend haplotypes on such “paths of length 1”, then the hypothesized reduction of 1/3 is achieved for long sequences. It is possible for extending haplotypes with Algorithm 1 to add edges, as in Figure 2, but we still expect the number of edges to decrease by more than 1/3. Indeed, Figure 4 shows that Algorithm 1 nearly cuts the number of edges in half, as long as the sequence is long enough.

**Figure 4.**
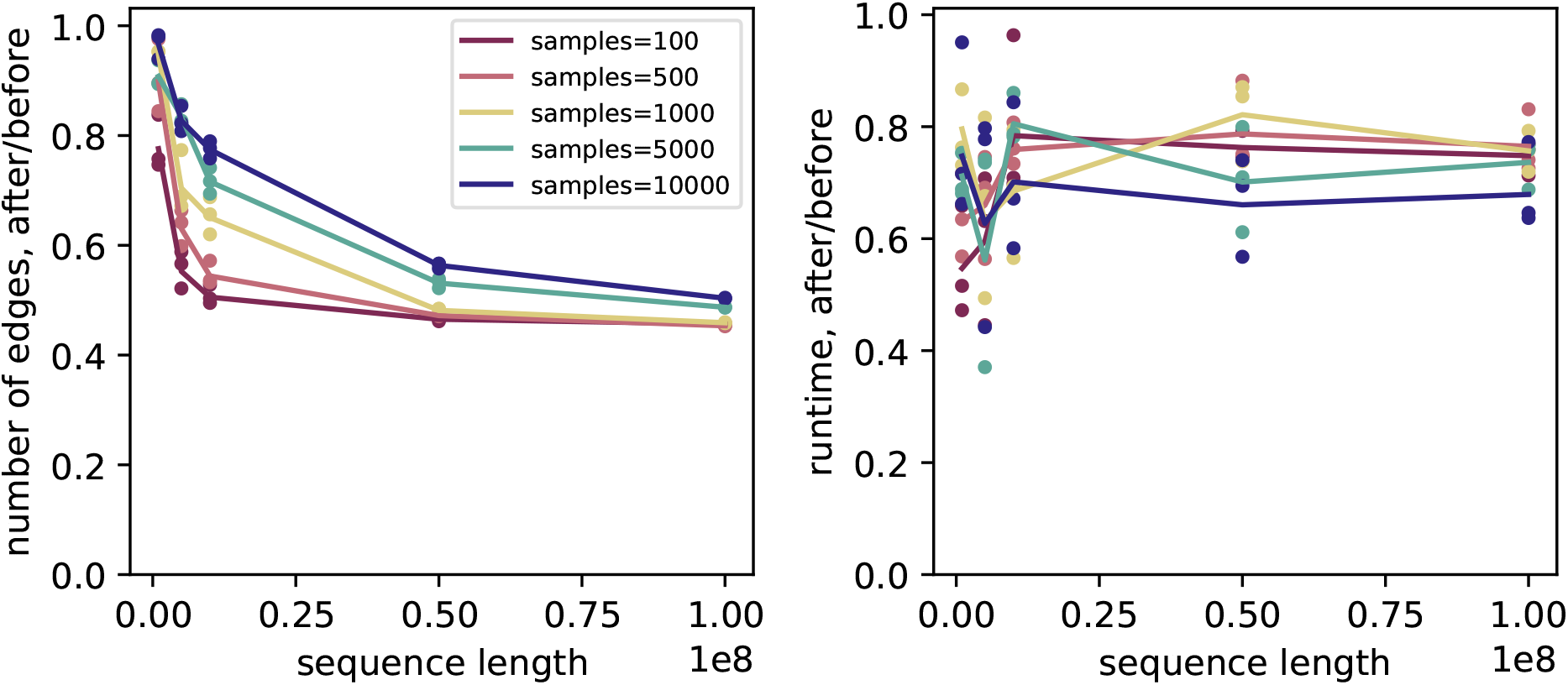
Ratio of **(A)** number of edges, and **(B)** runtime for computing Tajima’s *D*; before and after extending haplotypes. For instance, extending haplotypes reduces number of edges by about 50% and statistic computation runtime by about 20% for long sequences. Horizontal axis shows sequence length; colors show numbers of samples; with lines showing averages across replicates. The original tree sequence was simulated with the “expanding dog” expanding population and subset to various sizes; see Methods for details. Absolute values are shown in Supplementary Figure S1.

This reduction in edges can also lead to a reduction in computation time for algorithms using the succinct tree sequence data structure. Indeed, Figure 4 shows that computation time is reduced by 10–20% for a typical statistic (here, Tajima’s *D*), computed in an efficient incremental manner along the genome as implemented in `tskit`. As described in Ralph et al. [2020], for these incremental algorithms the addition or removal of an edge requires updates to the state of the parent node and all nodes ancestral to it. Extending haplotypes yields a tree sequence with fewer edge removals and insertions, and thus requires fewer traversals to the roots.

Supplementary Figure S2 shows these results are not specific to the demographic scenario. Supplementary Figures S1 and S3 also show that our implementation of Algorithm 1 is quite efficient, running at chromosome scale in seconds to minutes for hundreds or thousands of samples, or minutes to hours for tens of thousands of samples.

### 3.2 Accuracy with true trees

Our next task is to confirm that the haplotypes extended by Algorithm 1 are indeed correct – i.e., that in addition to compression, we are also gaining information. To do this, we simulate ARGs containing full haplotypes using Hudson’s algorithm in `msprime`, apply the simplification algorithm [Kelleher et al., 2018,Wong et al., 2024] to reduce these so that there are no unary nodes (i.e., any node present in a local tree is a coalescent node or a sample), and then apply Algorithm 1 to the result (see Methods for more detail). The method can potentially extend the spans of each node (additional span over which the node will be unary); and we can quantify how much of these extended spans were in the original ARG (and thus correctly extended).

As seen in Figure 5, the vast majority of span added by extending haplotypes is correct. In this example (which is typical), 99% of all added span is correct; 95% of nodes have no incorrectly added span; and those incorrectly added spans are nearly always a small fraction of the original span (Figure 5 B shows that incorrectly added span is often a factor of 10–100 smaller than the added span). The added information is significant: the algorithm typically increases spans (i.e., lengths of ancestral haplotypes) by around 50%. For instance, Figure 5 A shows that tens of megabases have been added to the spans of many nodes; while comparing Figure 5 C to Figure 5 D shows that much of the span removed by simplification has been replaced – in those figures, each point is one node; span after extending (points in D) are much closer to the original full span (solid blue points forming a line) than is span after simplifying (points in C). Some of the incorrectly added spans may be because of violations of the parsimony assumption, while others are due to situations with ambiguous information (and hence more than one way to extend a haplotype). It may be possible to improve the behavior of the algorithm in the latter situation, at the cost of a more complex algorithm.

**Figure 5.**
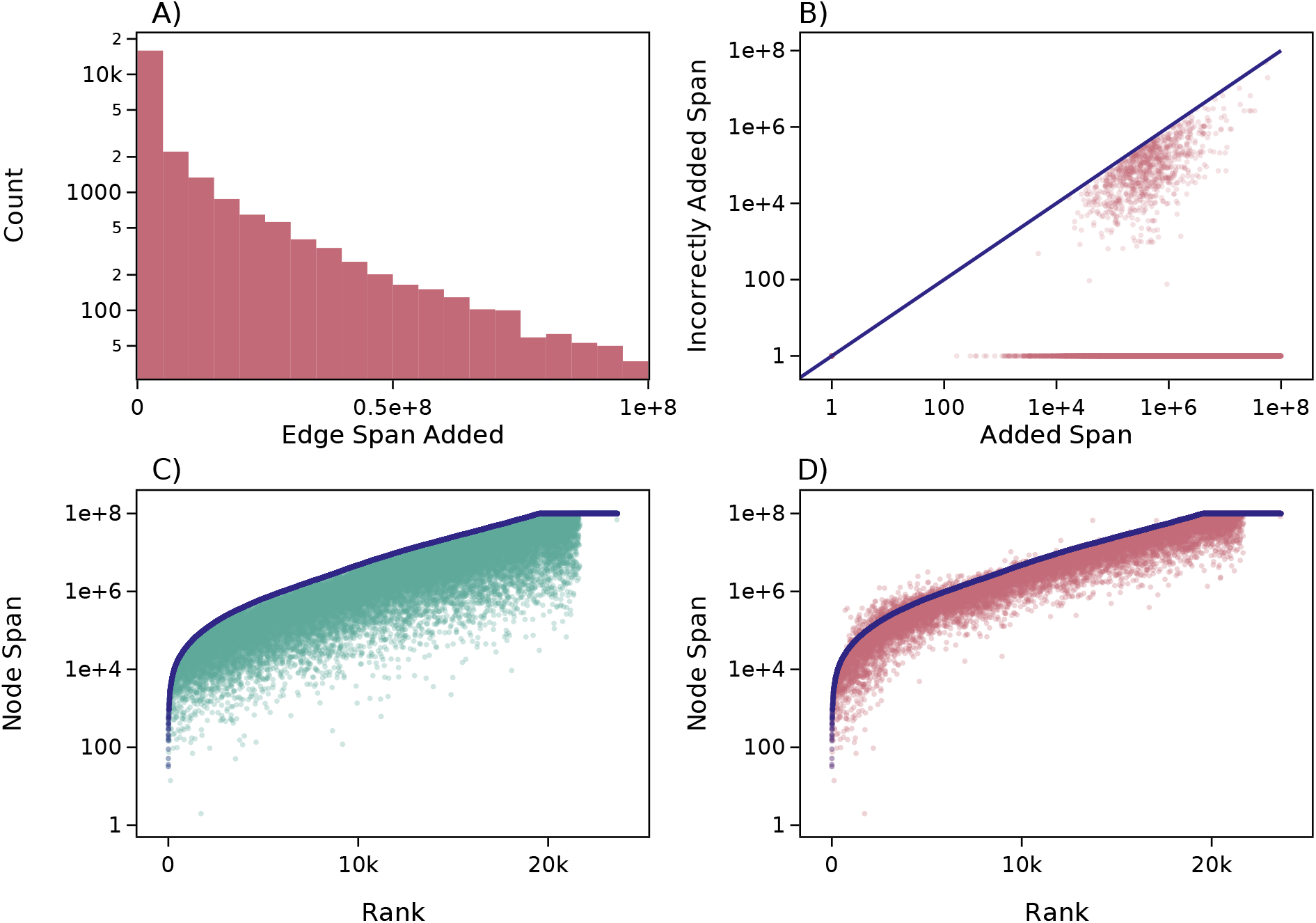
The effect of extending haplotypes on per-node spans in an ARG simulated with 10^4^ diploid samples in a population with *N*_*e*_ = 10^4^, and recombination rate of 10^™8^ on a sequence of length 108. **(A)** Distribution of total amount of span added across nodes by extending haplotypes with Algorithm 1; note the log scale on the *y* axis. **(B)** Amount of incorrectly added span, plotted against total span, by node. 95% of nodes have no incorrect span; of the remainder, nearly all have less than 5% incorrectly added; see Supplementary Figure S4. Note the log scale on both the *x* and *y* axes. Plots **(C)** and **(D)** show total spans per node, ordered by total span in the original ARG (which includes unary nodes). Dark blue dots in both show the span of each node in this original ARG. In **(C)**, lighter green dots show node span after removing unary spans using simplify, while in **(D)**, lighter red dots show node span after extending haplotypes, i.e., applying Algorithm 1 to the simplified ARG.

These statistics are also reflected at the genomic scale, using measures of matched span. Line “SE” in Figure 6 shows total amounts of span removed by simplification and re-inferred by extending haplotypes with Algorithm 1 (correctly and incorrectly). (Lines labeled with “I” involve re-inference of the ARG; discussed next.) The top row shows the proportion of the given ARG that does not match the original (“ARF”), showing that the total amount of mis-matching span produced by extending haplotypes is very small (≈ 1%). (Simplification does not produce non-matched span, so the “S” line is at zero.) The bottom row shows the proportion of the original ARG that is represented in the given ARG (“TPR”). This shows that a large proportion of an ARG can be removed by simplification, indicating that coalescent nodes are unary over a substantial portion of their spans. (Since the `simplify` operations removes these unary portions of haplotypes, the “simplified” line (“S”) on the bottom two plots of Figure 6 shows the proportion of nodes’ spans on which they are not unary.) However, Algorithm 1 can correctly replace most of these portions of haplotypes, especially with larger sample sizes and sequence lengths (the “simplified–extended” line; “SE”). For instance, the rightmost points show that with 1,000 samples and a 5 × 10^7^bp genome, haplotypes are unary on about half their spans (on average), and extending haplotypes can infer more than half of this missing unary span from coalescent information only.

**Figure 6.**
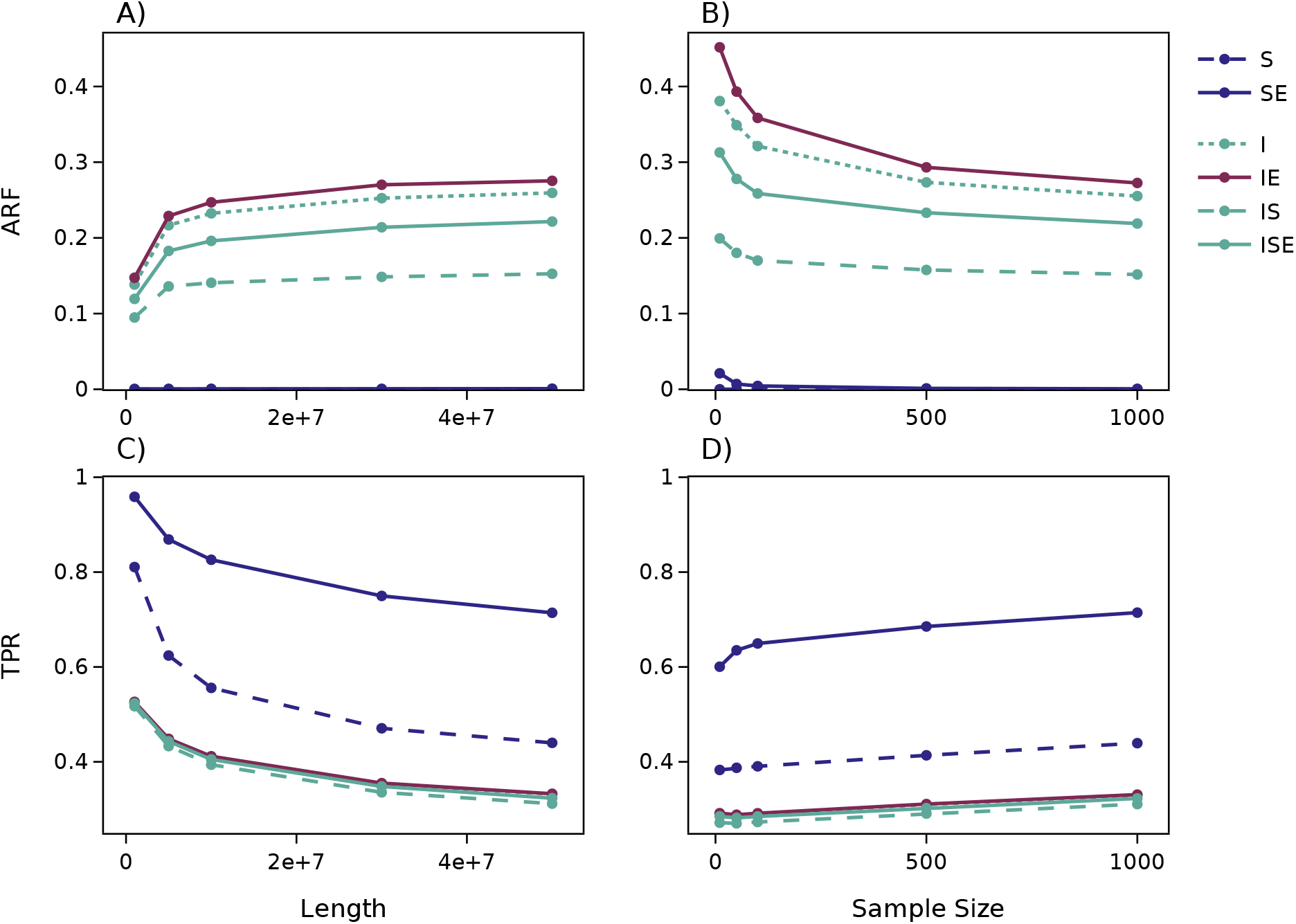
Accuracy and sensitivity of extended haplotypes across a range of sample sizes, on a sequence of length 5 *×* 10^7^ (right); and a range of sequence lengths, with 1,000 diploid samples (left). For each, a simulated ARG containing unary haplotype spans was (i) *simplified* (‘S’), removing the unary spans, and (ii) *inferred* (‘I’), using `tsinfer` on genotypes and dated using `tsdate`; then each of these had its haplotypes *extended* (‘SE’, ‘IE’). The inferred, then simplified, ARG (‘IS’) and its subsequent extension (‘ISE’) are also shown. **Top row:** ARF, the dissimilarity to the true ARG, as proportion of haplotypes that are not represented in the true ARG (Equation 4). **Bottom row:** TPR, agreement to the true ARG as proportion of the true ARG that is represented.

While our results are accurate at both the individual and genomic scale, we suspect that corrections can be made to further increase accuracy. Algorithm 1 relies on only the structure of the ARG as a tree sequence (its nodes, edges, and local trees), and does not use any other parameters for inference. However, it is possible that information including the recombination rate or gene conversion rate of an ARG could improve accuracy and reduce the amount of incorrectly added span for individuals. We decided to omit incorporating this information for simplicity and efficiency of our construction.

### 3.3 Inferred ARGs

So far, we have demonstrated that there is potentially ample information in the coalescent-only trees to extend haplotypes. Does this work with *inferred* ARGs? As illustrated by Wong et al. [2024], there is a significant diversity in the structures inferred by current methods, and here we focus on `tsinfer` Kelleher et al. [2019] which infers ARGs containing unary nodes as a byproduct of its inference algorithm. A comparison across other ARG inference methods is left for future work. We simulated ARGs containing unary-spanning haplotypes (as above), then re-inferred ARGs from the associated genotypes and performed various operations on the results. Since our matched spans method breaks ties with time, we date the re-inferred ARG using `tsdate`. First, Figure 6 shows that `tsinfer` has a substantial portion of already–extended haplotypes: comparing the “inferred” (“I”; dotted green) line to the “inferred-simplified” (“IS”; dashed green) line we see that inclusion of these unary spans increases the amount of correctly inferred material by around 4% (bottom panels), but that roughly 20% of the unary spans in the inferred ARG are incorrect (top panels). Furthermore, comparing to the inferred-extend(“IE”; red) line, we see that extending the tree sequence out-put with tsinfer adds relatively little span (less than about 1%). Extending haplotypes (with Algorithm 1) of the inferred-and-then-simplified ARG (“ISE”; solid green line) produces an ARG with both less correct and incorrect span. Additionally, the “IE” and “ISE” ARGs contain approximately the same number of edges (difference of ≈ 1%). It is also helpful to note that accuracy (i.e., proportion of the true ARG that is inferred; bottom panels) greatly increases with larger sample sizes, possibly due to resolution of polytomies. (Recall that due to computational constraints, “correct” and “incorrect” spans are determined here by *exact* match of subtending samples.) We additionally provide values of “I”, “IS”, “IE”, and “ISE” in Table S2, and numbers of edges and runtime for Tajima’s D in Figure S5.

## 4 Data availability

The method to extend haplotypes described here is available through the `tskit` python and C APIs (https://tskit.dev/tskit) as `extend_haplotypes`; methods to compare ARGs are implemented in the `tscompare` python package (https://tskit.dev/tscompare). Scripts used to produce the results in this paper are available at https://github.com/hfr1tz3/haplotypes-and-ancestral-recombination-graphs.

## 5 Discussion

We began this study with the observation that the simple transformation of Figure 2 would reduce the number of edges in the succinct tree sequence representation ARGs. This is essentially a recombination-based parsimony argument, and we have shown that this line of reasoning leads to ARGs that are substantially more compact and faster to operate on, and that contain more complete information about true ancestral relationships. These extended ancestral haplotypes manifest as unary nodes in the local trees. Although a number of ARG inference methods may be taking advantage of this information, it is our impression that this source of information is not widely appreciated. In fact, due to the field’s focus on local trees rather than haplotypes, we had to develop a haplotype-aware measure of (dis)agreement between ARGs in order to study the accuracy of the proposed algorithm.

There are good reasons to think that lengthening the spans of ancestral haplotypes could lead to substantial gains in accuracy of ARG inference. For instance, information about inferring the age of a particular mutation derives almost entirely from constraints at nearby, linked sites. Extending ancestral haplotypes from one site into neighboring regions conceptually allows information from those local trees to inform age inference at that site as well.

We have also explored the degree to which `tsinfer` already makes use of this information, and whether this algorithm can be used to improve inference. The results do not provide a clear ordering: for instance, although `tsinfer`-produced ARGs have a substantial portion of correctly inferred unary haplotypes, removing these with simplification decreases both ARF (i.e., proportion of “wrong” haplotypes) and TPR (the proportion of the truth that is correctly inferred). Extending haplotypes restores a large amount of this correctly inferred span, but also introduces incorrect spans. Presumably, this occurs because both correctly and incorrectly inferred haplotypes are extended. Further work is needed to determine how the balance of “true and false positive rates” affects downstream uses, and whether results would differ if the requirement that sets of subtended nodes match exactly was relaxed. The efficient computational tools we have implemented (in `tskit` and `tscompare`) should facilitate this exploration.

Another consideration is storage and computational efficiency: extending haplotypes reduces the number of edges, and thus filesize and (usually) runtime for computation. Figure 4 shows that the case is clear for simulated ARGs, substantially reducing both size and runtime. However, Figure S5 shows the situation is more complex for inferred ARGs: extending haplotypes on a `tsinfer`-produced ARG actually increases runtime somewhat, but simplifying-then-extending reduces runtime dramatically while keeping the number of edges roughly the same. More work is needed to understand how generalizable this is and what the source of these effects are.

### Ignorance and omission in an ARG

As motivation, we presented above a “historical” view of ARGs – i.e., that each aspect of an inferred ARG is intended to represent a portion of some particular historical genome (for instance, the MRCA of some set of sampled genomes). Furthermore, Figure 1 A&B implicitly takes the position that relationships *not* depicted in an ARG are implied to not exist. As discussed in Wong et al. [2024], an alternative interpretation of the ARG depicted in Figure 1 A&B would be that we have no information as to how node 2 inherited from node 4 on the right-hand interval, rather than saying that the line of transmission specifically did not pass through node 3. The “simplification” algorithm [Kelleher et al., 2018] and the Hudson algorithm for coalescent simulation [Hudson, 1983, Kelleher et al., 2016] each specifically discard information about any such “non-coalescent” portions of ancestral haplotypes; so for ARGs produced by these algorithms, the correct interpretation is that the omission of unary spans reflects a lack of knowledge. In this paper, we have shown that, for the most part, this missing information can be imputed.

### Parsimony

Much of the early work on ARG inference aimed to extract as much information as possible out of the small datasets of the time, and so, roughly speaking, integrated over possible ARGs with the goal of inferring higher-level parameters: mostly, scaled mutation rate and recombination rate (for instance, Hudson and Kaplan [1985], Griffiths and Marjoram [1996], Kuhner et al. [2000], Stephens and Donnelly [2000], Fearnhead and Donnelly [2001]). However, the space of possible ARGs for a given dataset is extremely large, and other work aimed to identify the minimum number of recombinations needed to explain a given dataset under the infinite alleles model of mutation [e.g., Hein, 1990, Myers and Griffiths, 2003, Song and Hein, 2005], which turns out to be NP-complete [Wang et al., 2001]. So, the field turned to more heuristic methods – for instance, Minichiello and Durbin [2006] used an algorithm to produce “plausible” ARGs (i.e., those that explained the data with few mutations and recombinations), and searched for associations with traits in the resulting ensemble of ARGs. (See Wong et al. [2024] for more historical discussion.) Our approach for extending haplotypes follows the same logic, that an ARG with fewer recombination events is more parsimonious, and thus more likely. For this reason, it will occasionally be wrong even if the trees are correct, although in practice this source of error is likely much smaller than error in tree inference itself.

### IBD in ARGs

The term “identity by descent” (IBD) is used to mean many different (but related) things, and length distributions of shared IBD segments can be used for inference of recent demographic history [for instance, Al-Asadi et al., 2019, Ringbauer et al., 2017, Browning and Browning, 2015, Yang et al., 2016, Silcocks et al., 2023]. A commonly-used definition in the context of a given ARG says that the two genomes share an IBD segment if each has inherited the segment from their common ancestor along a single path [e.g., Ralph and Coop, 2013]. Largely for computational reasons, this is the definition that is used in `tskit`‘s IBD-finding methods (by G. Tsambos in `tskit` [Kelleher et al., 2024]; see also Tsambos et al. [2023]). However, many simulation and inference methods produce ARGs as shown in Figure 1 A&B, in which inheritance of a single segment is represented by more than one edge. This means that the `ibd_segments` method of `tskit` will return shorter segments than it ought to. However, as we have shown above, our method of extending haplotypes will modify the tree sequence so that the inherited segment is represented by a single edge (as in Figure 1 C&D). So, if our method (`extend_haplotypes`) is applied before finding IBD segments (with `ibd _segments`), then the resulting segments should much more accurately represent the IBD segments (in the “path” sense used here) implied by the tree sequence. Whether the resulting segments better match those predicted by theory depends also on the quality of tree inference. Note also that Huang et al. [2024] and Guo et al. [2024] both provide methods for tree sequences to compute a different definition of IBD (segments on which the MRCA does not change), which is unaffected by this issue. Further work is needed to understand how accurately IBD segments are inferred by various ARG inference methods.

## 6 Acknowledgments

The authors would like to thank Dr. Yan Wong for his helpful comments and suggestions. JK acknowledges EPSRC (research grant EP/X024881/1) and the Robertson Foundation. JK and PR acknowledge support from the NIH NHGRI (research grant HG012473).

---

**Figure S1:**
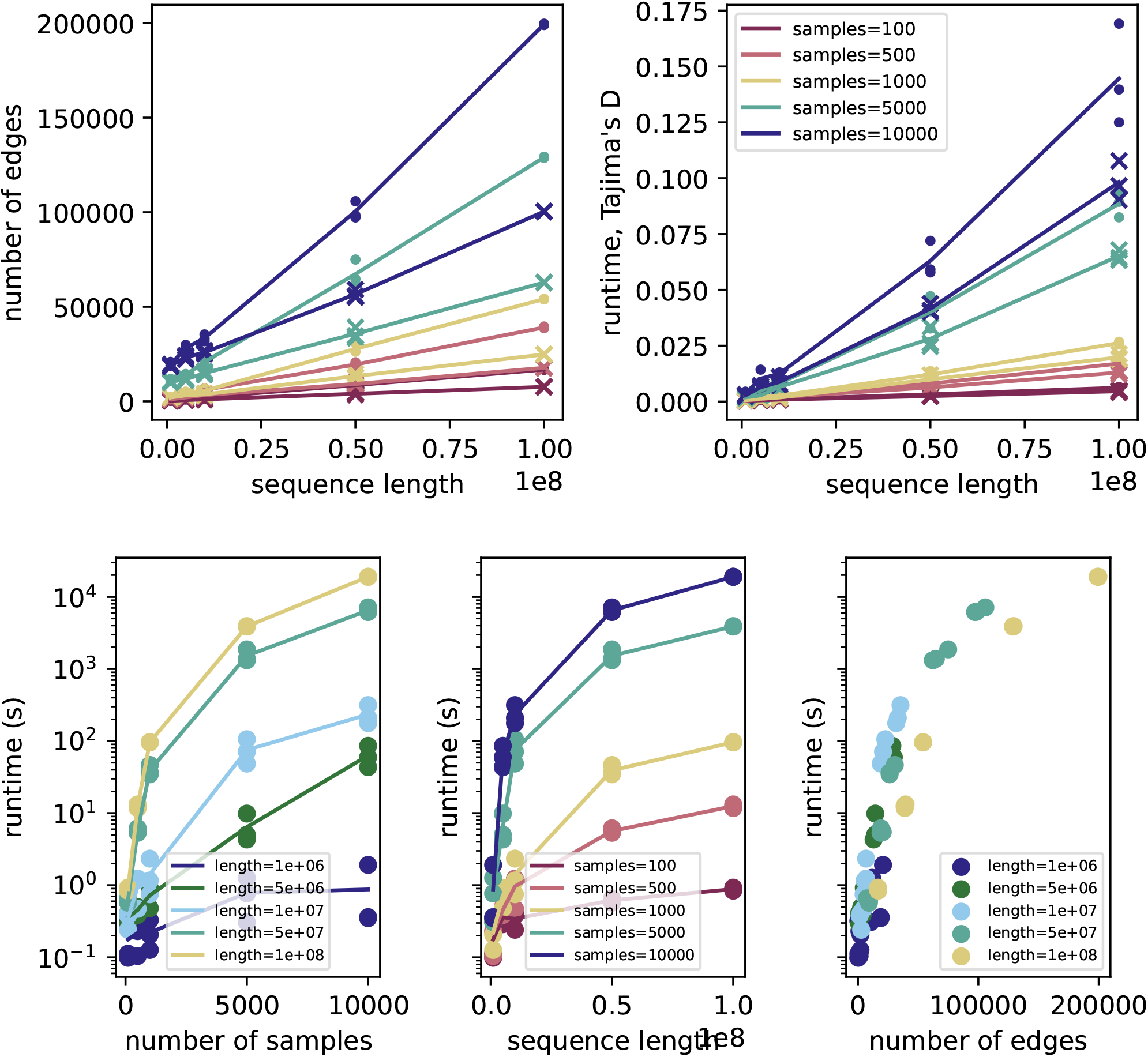
**(Above:)** Absolute values for ratios shown in Figure 4: **(left)** numbers of edges; and **(right)** runtime. **(Below:)** runtime for the `extend haplotypes` implementation of Algorithm 1 provided in the `tskit` library. For other details, see Figure 4.

**Figure S2:**
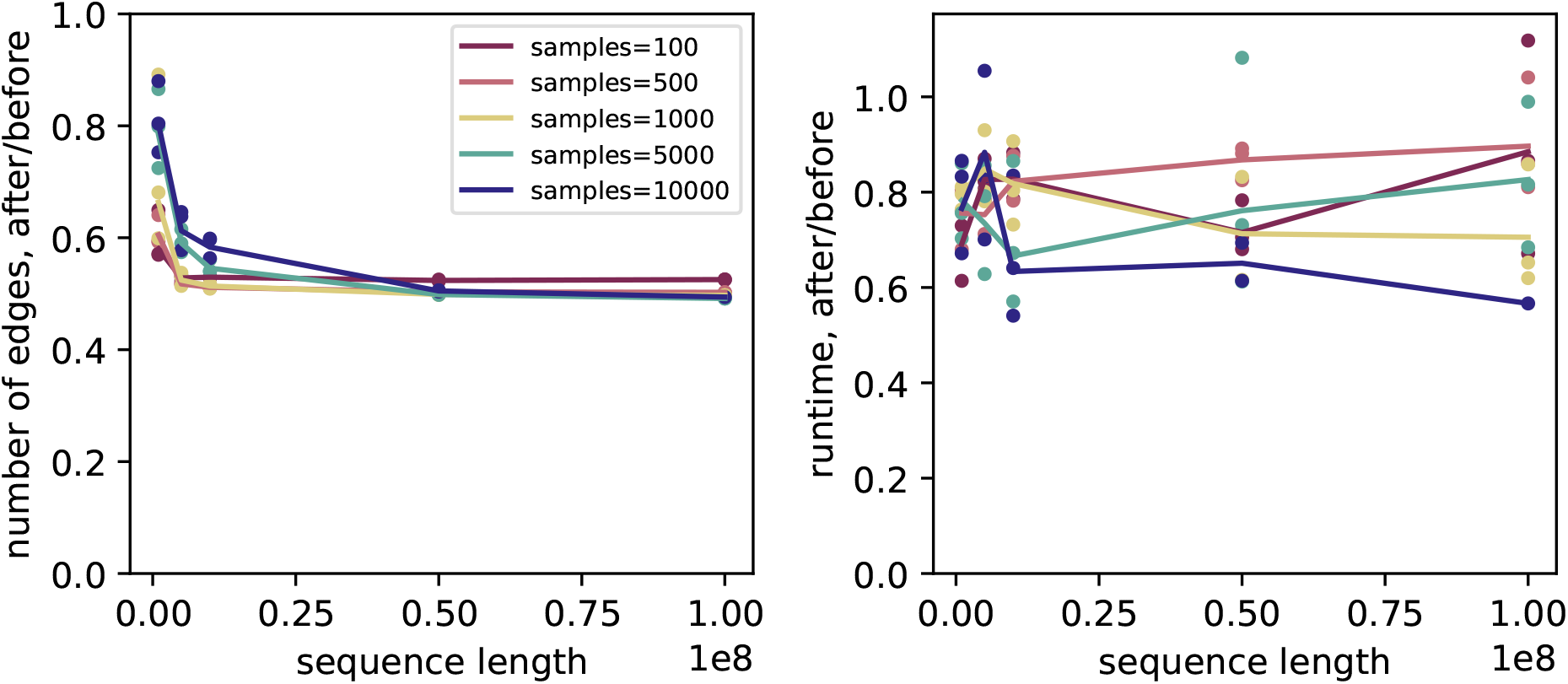
As in Figure 4 except that the original tree sequence was simulated with an constant population of size 10^4^ (using the “constant dog” scenario); see Methods for details. Absolute values are shown in Supplementary Figure S3.

**Table S1:** Number of edges for each iteration of *extend haplotypes* applied to simulated tree sequences with sample size 10^4^ and length 5 *×* 10^7^. Each of the simulations terminated by the fourth iteration, however 99% of the edges are removed within the first iteration.

| Initial | Iteration 1 | Iteration 2 | Iteration 3 | Iteration 4 |
| --- | --- | --- | --- | --- |
| 113463 | 76301 | 76059 | 76057 | 76057 |
| 113266 | 76084 | 75835 | 75833 | 75833 |
| 114489 | 76703 | 76436 | 76434 | 76434 |
| 114086 | 76550 | 76294 | 76291 | 76291 |

**Table S2:**
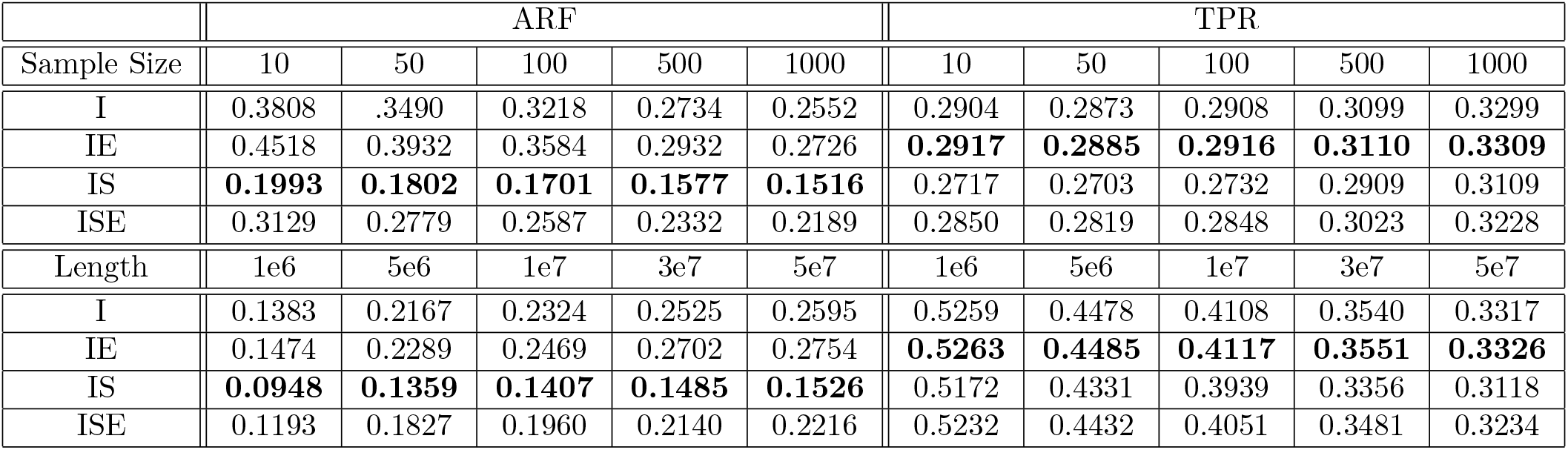
Values for ‘I’, ‘IE’, ‘IS’, ‘ISE’ in Figure 6.

**Figure S3:**
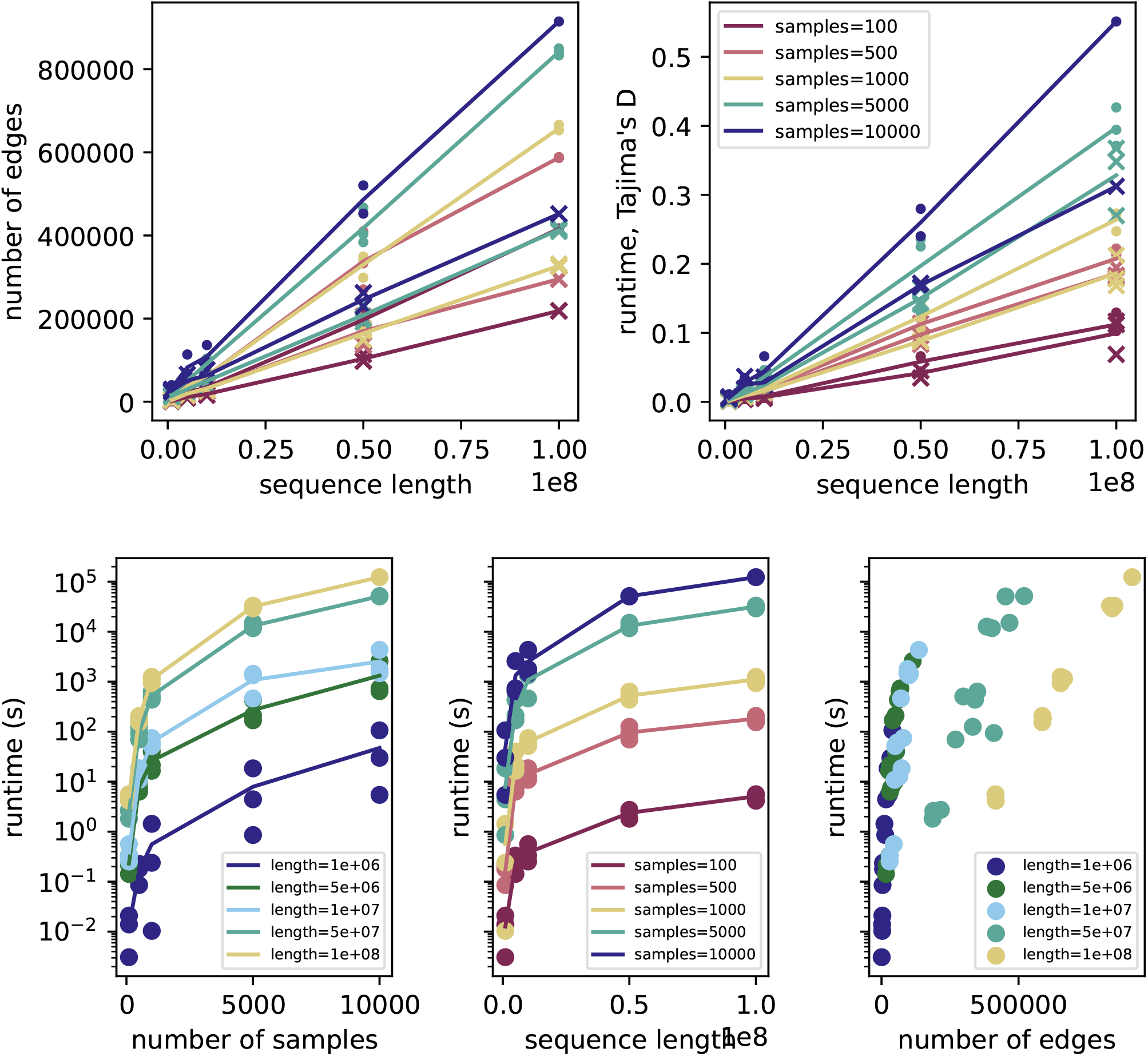
As in Figure S1 except that the original tree sequence was simulated with an constant population of size 10^4^ (using the “constant dog” scenario).

**Figure S4:**
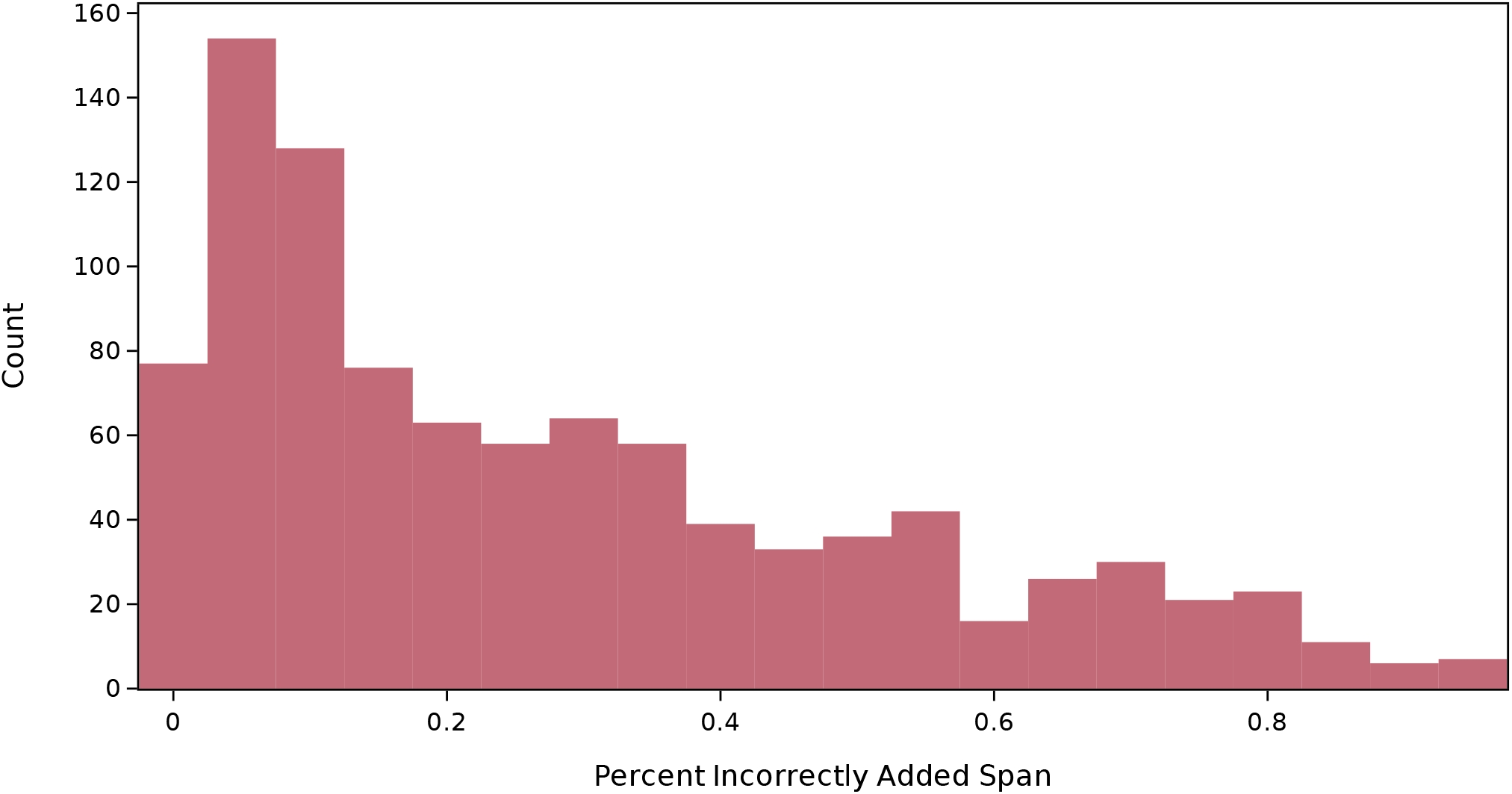
Distribution of incorrectly added span percentages for the simulated tree sequence in Figure 5.

**Figure S5:**
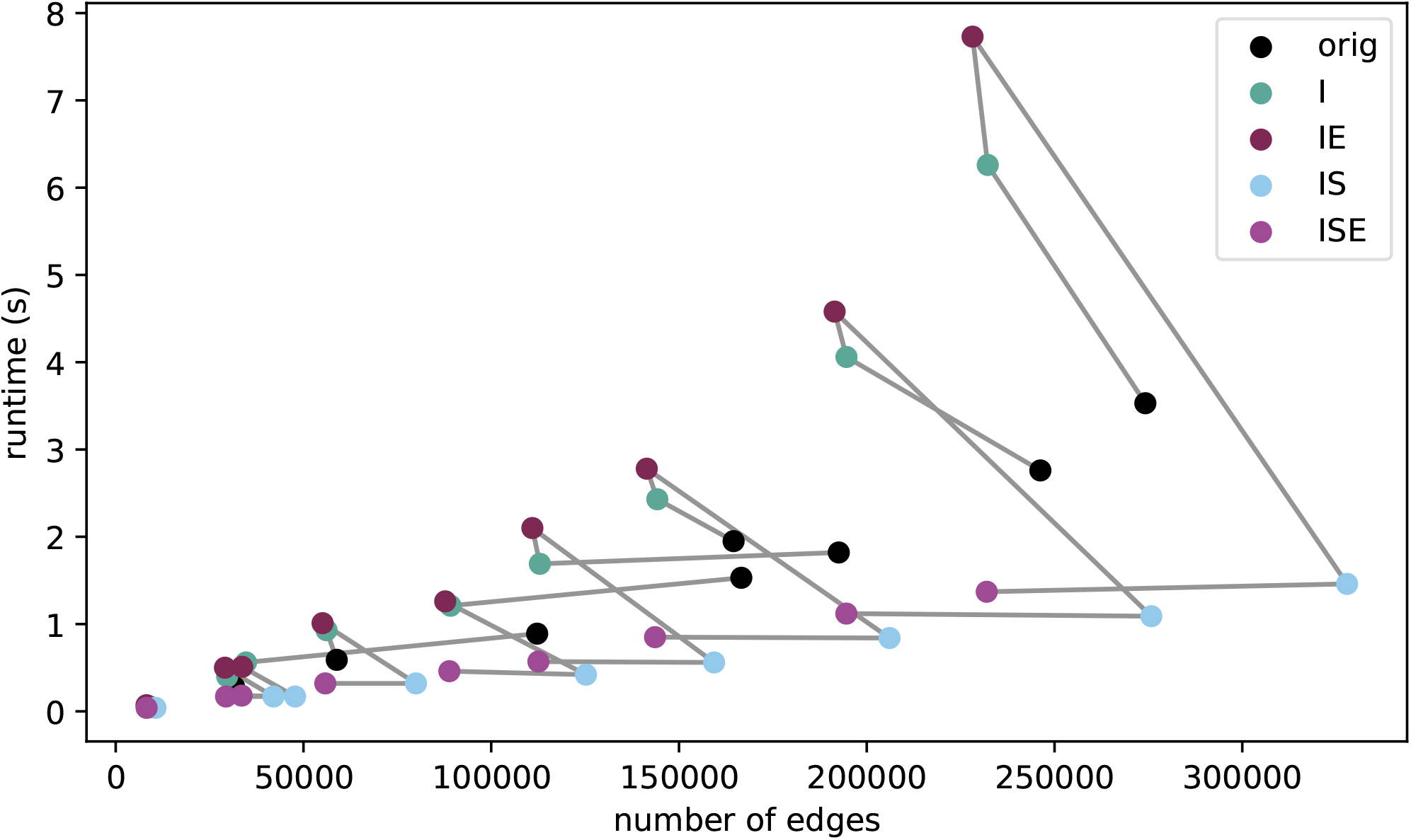
Numbers of edges and runtimes for computing Tajima’s D for the ARGs displayed in Figure 6. Grey lines connect each group of points that derive from the same original simulated ARG (i.e., each unique combination of sample size and sequence length). The label “orig” marks the originally simulated ARG.

## References

Jeffrey R Adrion, Christopher B Cole, Noah Dukler, Jared G Galloway, Ariella L Gladstein, Graham Gower, Christopher C Kyriazis, Aaron P Ragsdale, Georgia Tsambos, Franz Baumdicker, Jedidiah Carlson, Reed A Cartwright, Arun Durvasula, Ilan Gronau, Bernard Y Kim, Patrick McKenzie, Philipp W Messer, Ekaterina Noskova, Diego Ortega Del Vecchyo, Fernando Racimo, Travis J Struck, Simon Gravel, Ryan N Gutenkunst, Kirk E Lohmueller, Peter L Ralph, Daniel R Schrider, Adam Siepel, Jerome Kelleher, and Andrew D Kern. A community-maintained standard library of population genetic models. eLife, 9, June 2020. doi: 10.7554/elife.54967. URL https://doi.org/10.7554%2Felife.54967.

Hussein Al-Asadi, Desislava Petkova, Matthew Stephens, and John Novembre. Estimating recent migration and population-size surfaces. PLOS Genetics, 15(1):1–21, 01 2019. doi: 10.1371/journal.pgen.1007908. URL https://doi.org/10.1371/journal.pgen.1007908.

Franz Baumdicker, Gertjan Bisschop, Daniel Goldstein, Graham Gower, Aaron P Ragsdale, Georgia Tsambos, Sha Zhu, Bjarki Eldon, E Castedo Ellerman, Jared G Galloway, Ariella L Gladstein, Gregor Gorjanc, Bing Guo, Ben Jeffery, Warren W Kretzschumar, Konrad Lohse, Michael Matschiner, Dominic Nelson, Nathaniel S Pope, Consuelo D Quinto-Cortés, Murillo F Rodrigues, Kumar Saunack, Thibaut Sellinger, Kevin Thornton, Hugo van Kemenade, Anthony W Wohns, Yan Wong, Simon Gravel, Andrew D Kern, Jere Koskela, Peter L Ralph, and Jerome Kelleher. Efficient ancestry and mutation simulation with msprime 1.0. Genetics, 220(3):iyab229, 12 2021. ISSN 1943-2631. doi: 10.1093/genetics/iyab229. URL https://doi.org/10.1093/genetics/iyab229.

Sebastian Böcker, Stefan Canzar, and Gunnar W. Klau. The generalized Robinson-Foulds metric. In Aaron Darling and Jens Stoye, editors, Algorithms in Bioinformatics, pages 156–169, Berlin, Heidelberg, 2013. Springer Berlin Heidelberg. ISBN 978-3-642-40453-5.

Débora Brandt, Xinzhu Wei, Yun Deng, Andrew H Vaughn, and Rasmus Nielsen. Evaluation of methods for estimating coalescence times using ancestral recombination graphs. Genetics, 221(1):iyac044, 2022.

Débora YC Brandt, Christian D Huber, Charleston WK Chiang, and Diego Ortega-Del Vecchyo. The promise of inferring the past using the ancestral recombination graph. Genome Biology and Evolution, 16 (2):evae005, 2024.

Sharon R. Browning and Brian L. Browning. Accurate non-parametric estimation of recent effective population size from segments of identity by descent. The American Journal of Human Genetics, 97(3):404 – 418, 2015. ISSN 0002-9297. doi: 10.1016/j.ajhg.2015.07.012. URL http://www.sciencedirect.com/science/article/pii/S0002929715002888.

Christopher L Campbell, Claude Bhérer, Bernice E Morrow, Adam R Boyko, and Adam Auton. A pedigree-based map of recombination in the domestic dog genome. G3 Genes—Genomes—Genetics, 6(11):3517– 3524, 11 2016. ISSN 2160-1836. doi: 10.1534/g3.116.034678. URL https://doi.org/10.1534/g3.116.034678.

N. H. Chapman and E. A. Thompson. A model for the length of tracts of identity by descent in finite random mating populations. Theor Popul Biol, 64:141–150, September 2003. URL http://www.ncbi.nlm.nih.gov/pubmed/12948676.

Yun Deng, Yun S Song, and Rasmus Nielsen. The distribution of waiting distances in ancestral recombination graphs. Theoretical population biology, 141:34–43, October 2021. ISSN 00405809. doi: 10.1016/j.tpb.2021.06.003. URL https://pubmed.ncbi.nlm.nih.gov/34186053.

Yun Deng, Rasmus Nielsen, and Yun S. Song. Robust and accurate Bayesian inference of genome-wide genealogies for large samples. bioRxiv, 2024. doi: 10.1101/2024.03.16.585351. URL https://www.biorxiv.org/content/early/2024/03/16/2024.03.16.585351.

Paul Fearnhead and Peter Donnelly. Estimating recombination rates from population genetic data. Genetics, 159(3):1299–1318, 11 2001. ISSN 1943-2631. doi: 10.1093/genetics/159.3.1299. URL https://doi.org/10.1093/genetics/159.3.1299.

Ronald A Fisher. A fuller theory of ‘junctions’ in inbreeding. Heredity, 8(2):187–197, August 1954. ISSN 0018067X. URL http://dx.doi.org/10.1038/hdy.1954.17.

R.C. Griffiths and P. Marjoram. Ancestral inference from samples of DNA sequences with recombination. Journal of Computational Biology, 3(4):479–502, 1996. doi: 10.1089/cmb.1996.3.479. URL https://doi.org/10.1089/cmb.1996.3.479. PMID: 9018600.

Árni Freyr Gunnarsson, Jiazheng Zhu, Brian C. Zhang, Zoi Tsangalidou, Alex Allmont, and Pier Francesco Palamara. A scalable approach for genome-wide inference of ancestral recombination graphs. bioRxiv, 2024. doi: 10.1101/2024.08.31.610248. URL https://www.biorxiv.org/content/early/2024/09/02/2024.08.31.610248.

Bing Guo, Victor Borda, Roland Laboulaye, Michele D. Spring, Mariusz Wojnarski, Brian A. Vesely, Joana C. Silva, Norman C. Waters, Timothy D. O’Connor, and Shannon Takala-Harrison. Strong positive selection biases identity-by-descent-based inferences of recent demography and population structure in Plasmodium falciparum. Nature Communications, 15(1):2499, March 2024. ISSN 2041-1723. doi: 10.1038/s41467-024-46659-0. URL https://doi.org/10.1038/s41467-024-46659-0.

Jotun Hein. Reconstructing evolution of sequences subject to recombination using parsimony. Mathematical Biosciences, 98(2):185–200, 1990. ISSN 0025-5564. doi: 10.1016/0025-5564(90)90123-G.URL https://www.sciencedirect.com/science/article/pii/002555649090123G.

Zhendong Huang, Jerome Kelleher, Yao-ban Chan, and David J. Balding. Estimating evolutionary and demographic parameters via ARG-derived IBD. bioRxiv, 2024. doi: 10.1101/2024.03.07.583855. URL https://www.biorxiv.org/content/early/2024/03/13/2024.03.07.583855.1.

RR Hudson and NL Kaplan. Statistical properties of the number of recombination events in the history of a sample of DNA sequences. Genetics, 111(1):147–164, September 1985. URL http://www.ncbi.nlm.nih.gov/pmc/articles/PMC1202594/.

Richard R. Hudson. Properties of a neutral allele model with intragenic recombination. Theoretical Population Biology, 23(2):183–201, 1983. ISSN 0040-5809. doi: 10.1016/0040-5809(83)90013-8. UR https://www.sciencedirect.com/science/article/pii/0040580983900138.

Anastasia Ignatieva, Martina Favero, Jere Koskela, Jaromir Sant, and Simon R. Myers. The length of haplotype blocks and signals of structural variation in reconstructed genealogies. bioRxiv, 2024. doi: 10.1101/2023.07.11.548567. URL https://www.biorxiv.org/content/early/2024/06/19/2023.07.11.548567.

Jerome Kelleher, Alison M Etheridge, and Gilean McVean. Efficient coalescent simulation and genealogical analysis for large sample sizes. PLoS computational biology, 12(5):e1004842, 2016.

Jerome Kelleher, Kevin R. Thornton, Jaime Ashander, and Peter L. Ralph. Efficient pedigree recording for fast population genetics simulation. PLOS Computational Biology, 14(11):1–21, 11 2018. doi: 10.1371/journal.pcbi.1006581. URL https://doi.org/10.1371/journal.pcbi.1006581.

Jerome Kelleher, Yan Wong, Anthony W. Wohns, Chaimaa Fadil, Patrick K. Albers, and Gil McVean. Inferring whole-genome histories in large population datasets. Nature Genetics, 51(9):1330–1338, 2019. ISSN 15461718. doi: 10.1038/s41588-019-0483-y. URL https://doi.org/10.1038/s41588-019-0483-y.

Jerome Kelleher, Ben Jeffery, Yan Wong, Peter Ralph, Georgia Tsambos, Shing Hei Zhan, Kevin R. Thornton, Daniel Goldstein, Anthony Wilder Wohns, Duncan Mbuli-Robertson, Lloyd Kirk, Graham Gower, Hugo van Kemenade, Murillo F. Rodrigues, Brian Zhang, Gertjan Bisschop, Clemens Weiss, Duncan Palmer, Castedo Ellerman, Jeremy Guez, Nate Pope, Savita Karthikeyan, Inés Rebollo, Saurabh Belsare, Andrew Kern, and Maz Aspbury. tskit-dev/tskit: Python 0.5.8, June 2024. URL 10.5281/zenodo.12570616.

Michelle Kendall and Caroline Colijn. Mapping phylogenetic trees to reveal distinct patterns of evolution. Molecular Biology and Evolution, 33(10):2735–2743, 06 2016. ISSN 0737-4038. doi: 10.1093/molbev/msw124. URL https://doi.org/10.1093/molbev/msw124.

MK Kuhner and J Felsenstein. A simulation comparison of phylogeny algorithms under equal and unequal evolutionary rates. Molecular Biology and Evolution, 11(3):459–468, May 1994. ISSN 07374038. doi: 10.1093/oxfordjournals.molbev.a040126. URL https://pubmed.ncbi.nlm.nih.gov/8015439.

Mary K Kuhner and Jon Yamato. Assessing differences between ancestral recombination graphs. Journal of Molecular Evolution, 80(5):258–264, 2015.

Mary K Kuhner, Jon Yamato, and Joseph Felsenstein. Maximum likelihood estimation of recombination rates from population data. Genetics, 156(3):1393–1401, 11 2000. ISSN 1943-2631. doi: 10.1093/genetics/156.3.1393. URL https://doi.org/10.1093/genetics/156.3.1393.

Alexander L. Lewanski, Michael C. Grundler, and Gideon S. Bradburd. The era of the ARG: An introduction to ancestral recombination graphs and their significance in empirical evolutionary genomics. PLOS Genetics, 20(1):1–24, 01 2024. doi: 10.1371/journal.pgen.1011110. URL https://doi.org/10.1371/journal.pgen.1011110.

Mercé Llabrés, Francesc Rosselló, and Gabriel Valiente. The generalized Robinson-Foulds distance for phylogenetic trees. Journal of Computational Biology, 28(12):1181–1195, 2021. doi: 10.1089/cmb.2021.0342. URL https://doi.org/10.1089/cmb.2021.0342. PMID: 34714118.

Mark J. Minichiello and Richard Durbin. Mapping trait loci by use of inferred ancestral recombination graphs. The American Journal of Human Genetics, 79(5):910 – 922, 2006. ISSN 0002-9297. doi: 10.1086/508901. URL http://www.sciencedirect.com/science/article/pii/S0002929707608349.

Simon R Myers and Robert C Griffiths. Bounds on the minimum number of recombination events in a sample history. Genetics, 163(1):375–394, January 2003. ISSN 1943-2631. doi: 10.1093/genetics/163.1.375. URL http://dx.doi.org/10.1093/genetics/163.1.375.

Rasmus Nielsen, Andrew H Vaughn, and Yun Deng. Inference and applications of ancestral recombination graphs. Nature Reviews Genetics, pages 1–12, 2024.

Peter Ralph and Graham Coop. The geography of recent genetic ancestry across Europe. PLoS Biol, 11(5): e1001555, 05 2013. doi: 10.1371/journal.pbio.1001555. URL http://dx.doi.org/10.1371%2Fjournal.pbio.1001555.

Peter Ralph, Kevin Thornton, and Jerome Kelleher. Efficiently summarizing relationships in large samples: A general duality between statistics of genealogies and genomes. Genetics, page genetics.303253.2020, May 2020. doi: 10.1534/genetics.120.303253. URL https://doi.org/10.1534%2Fgenetics.120.303253.

Matthew D Rasmussen, Melissa J Hubisz, Ilan Gronau, and Adam Siepel. Genome-wide inference of ancestral recombination graphs. PLOS Genetics, 10(5):e1004342, 2014.

H Ringbauer, G Coop, and NH Barton. Inferring recent demography from isolation by distance of long shared sequence blocks. Genetics, 205(3):1335–1351, March 2017. doi: 10.1534/genetics.116.196220. URL https://www.ncbi.nlm.nih.gov/pubmed/28108588.

D. F. Robinson and L. R. Foulds. Comparison of weighted labelled trees. In A. F. Horadam and W. D. Wallis, editors, Combinatorial Mathematics VI, pages 119–126, Berlin, Heidelberg, 1979. Springer Berlin Heidelberg. ISBN 978-3-540-34857-3.

D.F. Robinson and L.R. Foulds. Comparison of phylogenetic trees. Mathematical Biosciences, 53(1):131– 147, 1981. ISSN 0025-5564. doi: 10.1016/0025-5564(81)90043-2. URL https://www.sciencedirect.com/science/article/pii/0025556481900432.

Matthew Silcocks, Ashley Farlow, Azure Hermes, Georgia Tsambos, Hardip R. Patel, Sharon Huebner, Gareth Baynam, Misty R. Jenkins, Damjan Vukcevic, Simon Easteal, Stephen Leslie, The National Centre for Indigenous Genomics, Ashley Farlow, Azure Hermes, Hardip R. Patel, Sharon Huebner, Gareth Baynam, Misty R. Jenkins, Simon Easteal, and Stephen Leslie. Indigenous Australian genomes show deep structure and rich novel variation. Nature, 624(7992):593–601, December 2023. ISSN 1476-4687. doi: 10.1038/s41586-023-06831-w. URL https://doi.org/10.1038/s41586-023-06831-w.

Yun S. Song and Jotun Hein. Constructing minimal ancestral recombination graphs. Journal of Computational Biology, 12(2):147–169, 2005. doi: 10.1089/cmb.2005.12.147. URL https://doi.org/10.1089/cmb.2005.12.147.

Leo Speidel, Marie Forest, Sinan Shi, and Simon R. Myers. A method for genome-wide genealogy estimation for thousands of samples. Nature Genetics, 51(9):1321–1329, 2019. ISSN 15461718. doi: 10.1038/s41588-019-0484-x. URL https://doi.org/10.1038/s41588-019-0484-x.

Matthew Stephens and Peter Donnelly. Inference in molecular population genetics. Journal of the Royal Statistical Society: Series B (Statistical Methodology), 62(4):605–635, 2000. doi: 10.1111/1467-9868.00254. URL https://rss.onlinelibrary.wiley.com/doi/abs/10.1111/1467-9868.00254.

Georgia Tsambos, Jerome Kelleher, Peter Ralph, Stephen Leslie, and Damjan Vukcevic. link-ancestors: fast simulation of local ancestry with tree sequence software. Bioinformatics Advances, 3(1):vbad163, 11 2023. ISSN 2635-0041. doi: 10.1093/bioadv/vbad163. URL https://doi.org/10.1093/bioadv/vbad163.

L Wang, K Zhang, and L Zhang. Perfect phylogenetic networks with recombination. Journal of computational biology : a journal of computational molecular cell biology, 8(1):69–78, 2001. ISSN 10665277. doi: 10.1089/106652701300099119. URL https://pubmed.ncbi.nlm.nih.gov/11339907.

Anthony Wilder Wohns, Yan Wong, Ben Jeffery, Ali Akbari, Swapan Mallick, Ron Pinhasi, Nick Patterson, David Reich, Jerome Kelleher, and Gil McVean. A unified genealogy of modern and ancient genomes. Science, 375(6583):eabi8264, 2022. doi: 10.1126/science.abi8264. URL https://www.science.org/doi/abs/10.1126/science.abi8264.

Yan Wong, Anastasia Ignatieva, Jere Koskela, Gregor Gorjanc, Anthony W Wohns, and Jerome Kelleher. A general and efficient representation of ancestral recombination graphs. Genetics, 228(1):iyae100, 07 2024. ISSN 1943-2631. doi: 10.1093/genetics/iyae100. URL https://doi.org/10.1093/genetics/iyae100.

Shuo Yang, Shai Carmi, and Itsik Pe’er. Rapidly registering identity-by-descent across ancestral recombination graphs. Journal of Computational Biology, 23(6):495–507, June 2016. ISSN 1557-8666. doi: 10.1089/cmb.2016.0016. URL http://dx.doi.org/10.1089/cmb.2016.0016.

Brian C. Zhang, Arjun Biddanda, Árni Freyr Gunnarsson, Fergus Cooper, and Pier Francesco Palamara. Biobank-scale inference of ancestral recombination graphs enables genealogical analysis of complex traits. Nature Genetics, 55(5):768–776, May 2023. ISSN 1546-1718. doi: 10.1038/s41588-023-01379-x. URL https://doi.org/10.1038/s41588-023-01379-x.

